# Integrated modeling of the Nexin-dynein regulatory complex reveals its regulatory mechanism

**DOI:** 10.1101/2023.05.31.543107

**Authors:** Avrin Ghanaeian, Sumita Majhi, Caitie L. McCaffrey, Babak Nami, Corbin S. Black, Shun Kai Yang, Thibault Legal, Ophelia Papoulas, Martyna Janowska, Melissa Valente-Paterno, Edward M. Marcotte, Dorota Wloga, Khanh Huy Bui

**Author notes:** Laboratory of Immunology, Mossakowski Institute of Experimental and Clinical Medicine, Polish Academy of Science, Pawinskiego 5, 02-106 Warsaw, Poland (Martyna Janowska).

## Abstract

Cilia are hairlike protrusions that project from the surface of eukaryotic cells and play key roles in cell signaling and motility. Ciliary motility is regulated by the conserved nexin-dynein regulatory complex (N-DRC), which links adjacent doublet microtubules and regulates and coordinates the activity of outer doublet complexes. Despite its critical role in cilia motility, the assembly and molecular basis of the regulatory mechanism are poorly understood. Here, utilizing cryo-electron microscopy in conjunction with biochemical cross-linking and integrative modeling, we localized 12 DRC subunits in the N-DRC structure of *Tetrahymena thermophila*. We also found that the CCDC96/113 complex is in close contact with the N-DRC. In addition, we revealed that the N-DRC is associated with a network of coiled-coil proteins that most likely mediates N-DRC regulatory activity.

## Introduction

Cilia are microscopic hair-like protrusions that extend from the surface of eukaryotic cells and are responsible for cell motility, sensory functions, and signaling. Ciliary motility has a critical role in the clearance of foreign particles from the respiratory system, establishment of the left-right asymmetry of the visceral body organs, development and maintenance of the spinal cord, development and composition of the brain ventricular system, and fertilization in the reproductive tract [1].

The motile cilium has a highly conserved core axial cytoskeleton called the axoneme. It is composed of a bundle of nine doublet microtubules (DMTs) composed of A- and B-tubules surrounded by two singlet microtubules known as the central pair (Fig. 1A). Each DMT is composed of multiple repeating 96-nm units comprising basically identical sets of axonemal complexes such as outer and inner dynein arms (ODAs and IDAs), radial spokes and a nexin-dynein regulatory complex (N-DRC) [2–5] (Fig. 1B). Dynein arms are A-tubule-docked molecular motors that generate a sliding force between DMTs [6–8]. ODAs generate high ciliary beat frequency, while IDAs control the bending shape of the cilia [9, 10]. Each 96-nm repeating unit contains four ODAs and seven IDAs, one two-headed IDA (dynein I1/f), and six distinct single-headed IDAs (dyneins a-g) with different mechanical properties [3]. The N-DRC bridges neighboring DMTs and restricts sliding motions between DMTs. As a result, the N-DRC converts dynein-generated sliding motion into ciliary bending motion.

**Figure 1.**
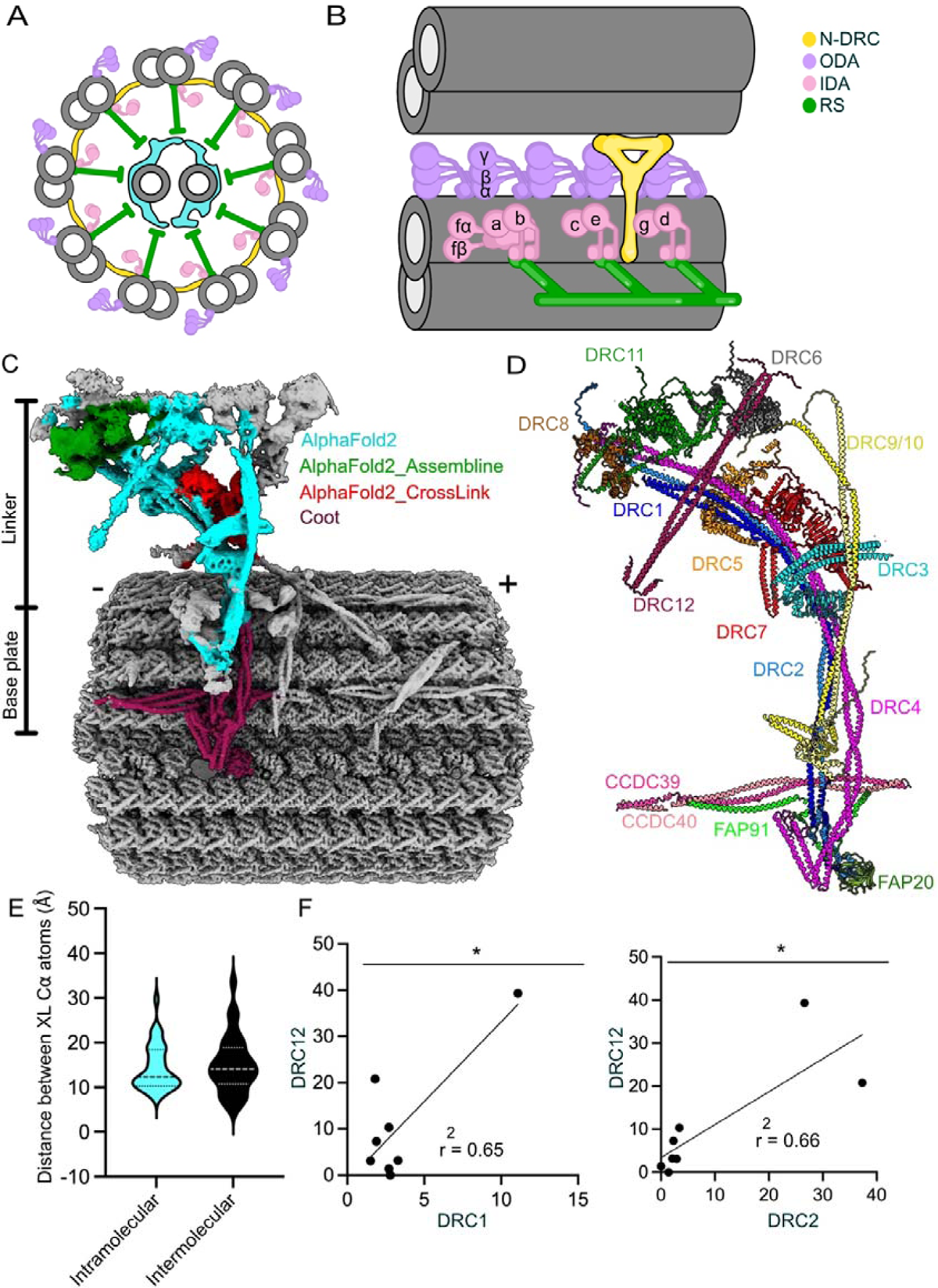
An overview of the structure of cilia and N-DRC. (A) Cross-sectional view of a cilium. (B) The 96-nm repeat of a DMT and its adjacent DMT is connected by the N-DRC (yellow) in longitudinal (left) and cross-sectional (right) views. (C) The composite cryo-EM map of N-DRC and DMT. The colors of different regions indicate the modeling methods. Signs (+) and (-) indicate the distal and proximal ends of the DMT. (D) The pseudo atomic model of the N-DRC. (E) Violin plots of the distance between inter-molecular and intra-molecular Cα atoms of chemically cross-linked residues. The intramolecular and intermolecular cross-links are shown in cyan and black, respectively. A maximum distance of 35 Å is expected for the DSSO crosslinks. (F) Correlation graphs of consensus normalized RNA expression levels for two selected pairs of genes (DRC12/DRC1 and DRC12/DRC2). The coefficient of the linear regression is indicated. *: *p*[<[0.05.

To generate the oscillating waveform movement of cilia, dynein activities switch back and forth from one side to the other side of the axoneme [11]. Such dynein regulation is accomplished through coordinated activities of mechano-regulatory complexes, including the central pair, radial spokes, N-DRC, and CCDC96/113 complexes [12] (Fig. 1B). Regulatory signals originating at the central pair are transmitted through the radial spokes and other mechano-regulatory complexes on the DMT to the dynein arms. The detailed mechanisms of signal transmission, however, remain unclear.

The N-DRC was first discovered in electron microscopy (EM) images of axonemal cross-sections and described as a nexin link that connects two adjacent DMTs [13]. Later, genetic studies revealed the presence of a “dynein regulatory complex” (DRC). Mutations in DRC subunit genes in Chlamydomonas rescued paralysis caused by mutations in radial spokes and central pair proteins [14–16]. Application of cryo-electron tomography (cryo-ET) finally revealed that the DRC and the nexin link are the same structure and coined the term “N-DRC” [17].

The N-DRC is a large 1.5-MDa, approximately 50-nm-long Y-shaped complex with two main regions: the base plate and linker regions (Fig. 1C). The base plate attaches to the inner junction of the DMT and four protofilaments of the A-tubule. The linker domain projects away through the inter-DMT space and connects the A-tubule to the neighboring B-tubule [17]. To date, biochemical and genetic studies have revealed that the N-DRC is composed of 11 subunits in *Chlamydomonas reinhardtii* and other species [18]. Cryo-ET experiments in wild-type and N-DRC mutants of *C. reinhardtii* revealed that DRC1, DRC2, and DRC4 span through the N-DRC from the base plate to the linker part, while DRC3, DRC5 to DRC11 were localized in the linker [17, 19–21]. A high-resolution cryo-electron microscopy (cryo-EM) study of the DMT from *Chlamydomonas* revealed that DRC1, DRC2, and DRC4 form the core structure of the base plate [22]. Mutations in DRC subunit genes of *C. reinhardtii* caused the disassembly of the N-DRC, defects in ciliary movement and destabilized the assembly of several closely associated structures, such as the IDAs [9, 23–25].

Despite extensive research on the N-DRC, the precise composition and structural model of the components have yet to be determined. Consequently, our understanding of the intra- and inter-interactions of the N-DRC and its function in dynein regulation and ciliary movement is still fragmentary. Here, we used integrated modeling and cryo-EM to model the entire N-DRC structure and some associated proteins from the ciliate *Tetrahymena thermophila*. Our study revealed the presence of 12 N-DRC components and their detailed organization. Moreover, our structural model also indicates that the N-DRC regulates ODAs and IDAs by interacting with a network of coiled-coil proteins.

## Results

### Integrated modeling reveals the molecular architecture of the N-DRC

To obtain the N-DRC structure, we performed single-particle analysis of isolated DMT from *T. thermophila* from several datasets (Table 1, Methods) [26, 27]. We obtained the 96-nm repeat of the DMT at a global resolution of 3.6 Å (Fig. S1), containing the N-DRC base plate. This allowed the detailed modeling of the N-DRC base plate, which consists of segments of DRC1, DRC2, DRC4A, DRC4B, and associated proteins on the surface of the DMT, CCDC39, CCDC40, and CFAP91 (Fig. S2A, Table. S1).

**Table 1:**
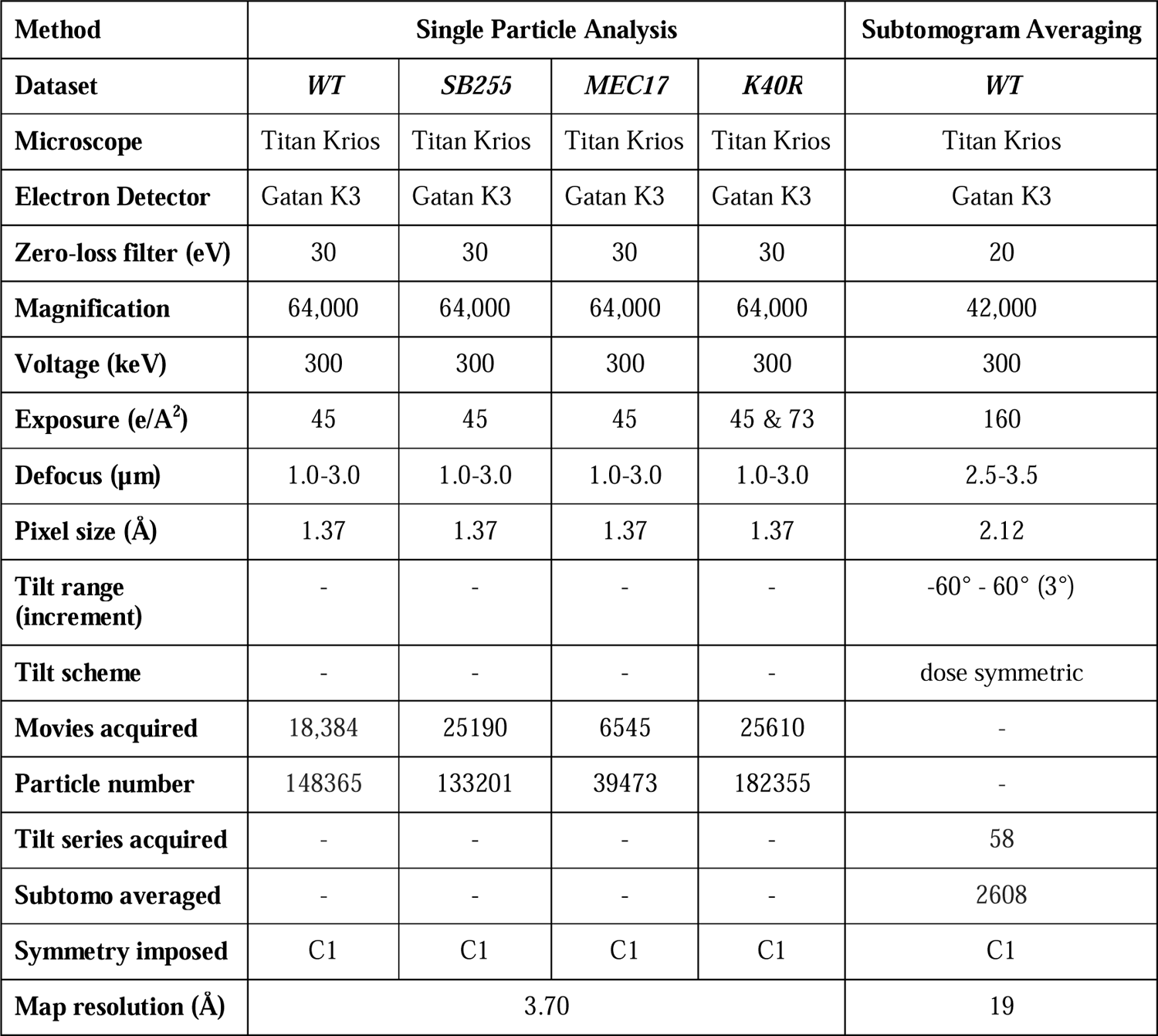
Cryo-EM data collection and refinement parameters for all datasets used in this study.

| Method | Single Particle Analysis |  |  |  | Subtomogram Averaging |
| --- | --- | --- | --- | --- | --- |
| Dataset | *WT* | *SB255* | *MEC17* | *K40R* | *WT* |
| Microscope | Titan Krios | Titan Krios | Titan Krios | Titan Krios | Titan Krios |
| Electron Detector | Gatan K3 | Gatan K3 | Gatan K3 | Gatan K3 | Gatan K3 |
| Zero-loss filter (eV) | 30 | 30 | 30 | 30 | 20 |
| Magnification | 64,000 | 64,000 | 64,000 | 64,000 | 42,000 |
| Voltage (keV) | 300 | 300 | 300 | 300 | 300 |
| Exposure (e/Å<sup>2</sup>) | 45 | 45 | 45 | 45 & 73 | 160 |
| Defocus (μm) | 1.0-3.0 | 1.0-3.0 | 1.0-3.0 | 1.0-3.0 | 2.5-3.5 |
| Pixel size (Å) | 1.37 | 1.37 | 1.37 | 1.37 | 2.12 |
| Tilt range (increment) | - | - | - | - | -60° - 60° (3°) |
| Tilt scheme | - | - | - | - | dose symmetric |
| Movies acquired | 18,384 | 25190 | 6545 | 25610 | - |
| Particle number | 148365 | 133201 | 39473 | 182355 | - |
| Tilt series acquired | - | - | - | - | 58 |
| Subtomo averaged | - | - | - | - | 2608 |
| Symmetry imposed | C1 | C1 | C1 | C1 | C1 |
| Map resolution (Å) | 3.70 |  |  |  | 19 |

**Table 2:** Refinement statistics of the base plate and linker models.

|  | **Base plate** | **Linker** |
| --- | --- | --- |
| **Model-to-Map fit, CCmask** | 0.43 | 0.59 |
| **All-atom clashscore** | 17.70 | 78.51 |
| **Ramachandran plot** |  |  |
| **Outliers [%]** | 0.11 | 0.04 |
| **Allowed [%]** | 3.45 | 5.72 |
| **Favored [%]** | 96.44 | 94.25 |
| **Rotamer outliers [%]** | 1.00 | 0.00 |
| **Cbeta deviations [%]** | 0.06 | 0.04 |
| **Cis-proline [%]** | 0.00 | 2.0 |
| **Cis-general [%]** | 0.1 | 0.1 |
| **Twisted proline [%]** | 3.7 | 1.3 |
| **Twisted general [%]** | 0.1 | 0.5 |

However, the N-DRC linker part was not well resolved due to its flexibility (Fig. S1). Therefore, we developed an image processing strategy to obtain the linker structure at ∼5-10 Å resolution involving extensive subtractions, three-dimensional classifications and focused refinement (Fig. S1, Methods). This resolution is sufficient to fit the AlphaFold2 predicted structures of N-DRC components unambiguously, especially the coiled-coil part of the proteins (Fig. 1C, Fig. S2B-F, Table S1-2, see Methods). As a result, we can confidently fit the remaining AlphaFold2 multimer model of DRC1/2, DRC4A/4B, DRC9/10 complexes, DRC3 and DRC5 into the N-DRC linker part (Fig. S2C-E).

For the remaining region where we could not fit in the predicted structures uniquely, we performed integrated modeling using the Assembline program and spatial restraints derived from cross-links identified by *in situ* cross-linking mass spectrometry of the *T. thermophila* cilia [28]. We localized DRC11A and two copies of DRC8 to the proximal lobe of the linker and DRC7 in the middle part of the linker (Fig. 1C, D). Integrated modeling yields two possible positions for DRC6A (Fig. S3A). Overall, our integrated model of the N-DRC and associated proteins explains ∼85% of the analyzed density (Fig. 1D, S2B). However, we could not localize proteins to the N-DRC distal lobe.

Next, we used the inter-molecular and intra-molecular and intra-molecular cross-links between N-DRC components and associated proteins from *in situ* cross-linking mass spectrometry to validate our model (Table S3, S4). Our model satisfies all the spatial restraints from crosslinks found for N-DRC proteins (Fig. 1E, S2B, and Table S3 and S4). Interestingly, we found a crosslink from a conserved protein, CCDC153 (UniProtID Q22RH5), to DRC1 in the linker region (Table S3). The density of the unknown coiled coil from the N-DRC linker region matches the AlphaFold2 predicted structure of the CCDC153 homodimer (Fig. 1C, D, and Fig. S4A). Accordingly, we propose that CCDC153 is an N-DRC subunit that we named DRC12 (Fig. 1C, D, S4A). To verify DRC12 localization, we performed BioID assays using DRC1-HA-BirA* and DRC2-HA-BirA* as baits. Most DRC components were biotinylated in both experiments, including DRC12 (Table S5), validating the proximity of DRC1 and DRC2 to DRC12. The high correlation of the normalized RNA expression of CCDC153 from different human tissues with that of DRC1 (CCDC164) and DRC2 (CCDC65) (Fig. 1F) and other N-DRC components (Fig. S4B) further supports the assignment of CCDC153 as a DRC protein.

DRC4 forms a homodimer in *C. reinhardtii* [22]. *T. thermophila* contains two DRC4 paralogs, DRC4A and DRC4B. The identified inter-molecular crosslinks between DRC4A and DRC4B but not intra-molecular crosslinks suggest that DRC4A and DRC4B predominantly form a heterodimer (Tables S3 and S4). To further support that DRC4A and DRC4B exist as heterodimers, we performed pull-down assays using anti-GFP beads and DRC4A-GFP or DRC4B-GFP as bait. Both proteins efficiently pulled down the HA-tagged DRC4 paralog, DRC4A-GFP pulled down DRC4B-HA and vice versa, DRC4B-GFP interacted with DRC4A-HA, while homodimeric interactions between DRC4 paralogs were poor or undetectable (Fig. S5A, B). Interestingly, the pulldown analysis indicated that DRC3-HA also interacted with DRC4B but poorly with DRC4A-GFP. Further analysis of the AlphaFold2 multimer prediction of DRC4A/4A, DRC4B/4B, and DRC4A/4B dimers shows that only the DRC4A/4B heterodimer matches the DRC4 density in our map (Fig. S5C, D). These results imply that in *T. thermophila,* the DRC4 dimer exists mainly as a DRC4A/4B heterodimer (Fig. S5C-D) (see Methods).

### The N-DRC core scaffold consists of coiled-coil proteins

The backbone structure of the N-DRC consists of coiled-coil proteins (DRC1, DRC2, DRC4A, DRC4B) that extend from the base plate to the linker part (Fig. 1D). These coiled-coil proteins can give the N-DRC elasticity.

In *T. thermophila* cilia, the C-terminal fragments of the DRC1/2 and DRC4A/4B heterodimers are located on the A-tubule and form the N-DRC base plate (Fig. 1D, S2A), similar to *C. reinhardtii* [22]. At the base plate, the DRC1/2 and DRC4A/4B heterodimers contact perpendicularly with the coiled-coil CCDC39/40, the so-called “molecular ruler” that runs along the DMT in the wedge of protofilaments A2A3 and determines the 96-nm repeat of the DMT. The C-terminal region of the DRC1/2 heterodimer also interacts with (i) the C-terminal region of the DRC4A/B heterodimer, (ii) CFAP91, which runs almost parallel to the CCDC39/40 coiled coil (Fig. 2A) and (iii) CFAP20, an inner junction protein [27, 29]. Previous studies showed that CFAP91 extends from the base of RS2 through the N-DRC base plate to the radial spoke RS3, playing a significant role in stabilizing and localizing radial spokes RS2 and RS3 on the DMT [22, 30]. N-DRC positioning on the DMT may depend on interactions between the DRC1/2 coiled coil and the DRC4A/B coiled coil and the CCDC39/40 coiled coil (Fig. 1C-D, 2A).

**Figure 2.**
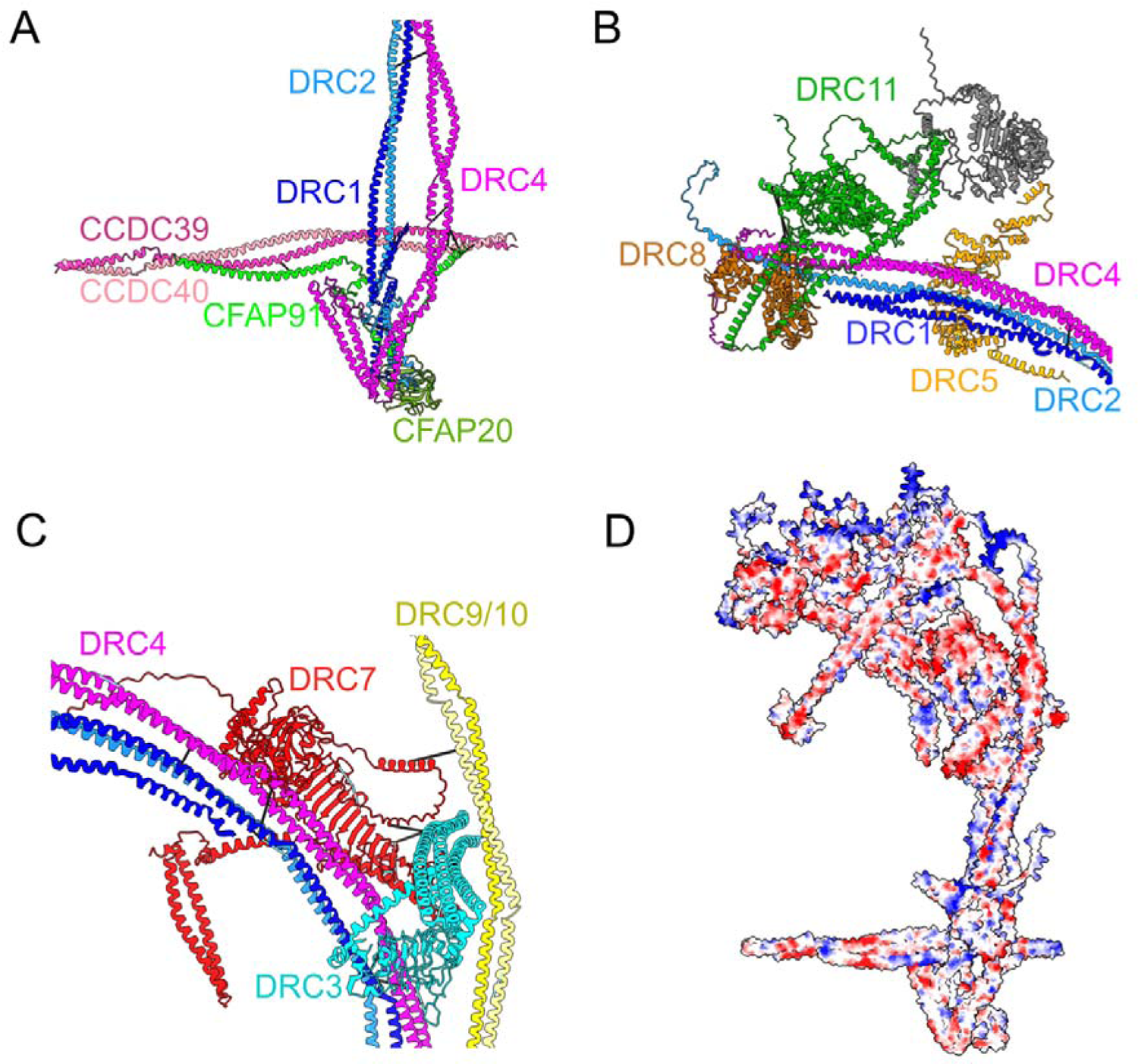
The interactions within the N-DRC. (A) The interactions in the base plate region. (B) DRC5 and DRC11/8 subcomplex interact with scaffold proteins (DRC1, DRC2, and DRC4). DRC12 is eliminated from the figure for clarity. (C) The interactions between proteins in the N-DRC distal lobe. DRC7 interacts with the DRC9/10 coiled coil and DRC3 in the linker distal part and with DRC4A/4B coiled coil in the proximal part. The cyan and black bars represent intramolecular and intermolecular cross-links, respectively. (D) The electrostatic surface charge of the N-DRC model. The loops at the top of the linker (DRC11A, DRC6A, DRC2, and DRC9/10), which connect to the adjacent DMT, are positively charged. The colors blue, red, and white indicate positive, negative and neutral charges, respectively.

The N-DRC core scaffold is composed of DRC1, DRC2 and DRC4, which interact with DRC3, DRC5, DRC8 and DRC11 in the linker part (Fig. 1D, 2B). Our model indicates that the DRC1/2 coiled coil interacts with the DRC4A/B coiled coil not only in the base plate region but also in the top part of the linker (Fig. 1D, 2B). This coiled-coil bundle constructs the N-DRC core structure (Fig. 1D). This suggests that the DRC1/2 and DRC4A/B complex is critical for the assembly of the N-DRC since they position the N-DRC base plate to the DMT via interaction with CFAP20 in the inner junction and interact with all the proteins in the N-DRC central and proximal lobes (Fig. 1D). This is consistent with the phenotypes of the *C. reinhardtii* mutants of *DRC1* (*pf3*) and *DRC2* (*ida6*), and human mutations in the *CCDC164* (*DRC1*) and *CCDC65* (*DRC2*) genes, which cause the complete loss of the N-DRC [24, 25, 31]. As an N-DRC core scaffold component, the loss of the DRC4 protein in the *pf2* mutant in *C. reinhardtii* causes the absence of all DRC subunits except the DRC1/2 complex [15, 18, 24].

DRC9/10 is another coiled-coil dimer that binds to the DRC1/2 coiled coil just above the base plate and projects through the linker part distally to form the other arm of the Y-shape of the N-DRC and interacts with DRC3 in the distal region (Fig. 1D, 2C).

Within the N-DRC linker region, the leucine-rich DRC5 subunit interacts extensively with DRC1/2 and DRC4A/4B coiled coils (Fig. 2B). In the top part of the linker, the extended N-terminus of DRC2 interacts with DRC8, while DRC1 folds back to the central part of the linker and interacts with DRC5 (Fig. 2B, S2C). The DRC4A/B heterodimer plays an essential role in the stability of the N-DRC by interacting with the leucine-rich domains of DRC3, DRC5, and DRC7 in the central part of the linker and interacting with DRC11 in the linker top part (Fig. 1D, 2B, C).

DRC7 is in the middle part of the linker, which connects the N-DRC proximal and distal lobes. In the N-DRC distal lobe, DRC7 interacts with the DRC9/10 coiled coil and the α-helix bundle domain of DRC3. In the proximal lobe, DRC7 interacts with DRC4A/4B coiled coil (Fig. 1D, 2C). Therefore, DRC7 can be a signal transfer bridge between the distal and proximal lobes. Although DRC7 connects the distal domain to the proximal lobes by linking DRC4A/B to DRC10, loss of DRC7 in the *drc7* mutant does not have any effect on the assembly of the distal subunits (DRC3, DRC9, and DRC10) [21], suggesting that DRC7 likely reinforces the distal lobe but is not essential for the assembly of the distal lobe.

From AlphaFold2 prediction and density fitting, our model predicted that two copies of EF-hand DRC8 proteins bind tightly to the helical hairpin domain of the AAA+ DRC11 protein (Fig. 2B), agreeing with the localization of DRC11 shown using *C. reinhardtii* [21] and *Trypanosome* [32] mutant strains. Interestingly, superimposition of the subtomogram average map of human DMT onto that of *T. thermophila* and AlphaFold2 Multimer modeling of the N-DRC in humans revealed that only one copy of DRC8 is present in the human N-DRC (Fig. S5E-F). DRC11 contains an AAA+ domain that potentially contributes to conformational changes in the N-DRC [18]. In addition, DRC11 has a regulatory role since the knockdown of DRC11 homolog (CMF22) in *trypanosomes* causes a motility defect [33].

The electrostatic surface charge of our N-DRC model indicates that the top of the linker part contains positively charged flexible loops (Fig. 2D). This allows the binding of the N-DRC to the negatively charged tubulin from the adjacent DMT [34]. This also serves as a validation for our N-DRC model since we are not using surface charge information while building it.

### The CCDC96/113 complex interacts with the N-DRC

In our cryo-EM map, the CCDC96/113 heterodimer, the conserved mechano-complex distal to the N-DRC [12], resolves well in the basal part, allowing *de novo* modeling of this region. The C-terminal ends of CCDC96 and CCDC113 form a clear coiled-coil domain (Fig. S6A). AlphaFold2 multimer prediction of CCDC96/113 is highly confident and accurately represents the topology of the coiled coil in the cryo-EM map, allowing unambiguous docking of the CCDC96/113 heterodimer in the region away from the DMT surface (Fig. S6A). The coiled-coil domain of CCDC96/113 is located parallel to the N-DRC, but from amino acid 458 onward, it forms a 90-degree turn toward the N-DRC distal part (Fig. 3A, S6B). The N-terminus of CCDC96 in *T. thermophila* is not conserved and contains an extra WD40 beta-propeller compared to humans and *C. reinhardtii* (Fig. S6C-D). Following the 90-degree turns of CCDC96, we predict that the non-conserved globular domain of CCDC96 extends to the N-DRC distal lobe by a flexible loop and localizes close to DRC10 (Fig. 3A). To support our prediction, we performed cryo-ET and subtomogram averaging to obtain the 96-nm repeat of *T. thermophila* at 20 Å resolution and compared it to that of *T. thermophila* CCDC96-deletion, *C. reinhardtii* (EMD-20338), and humans (EMD-5950). Apparently, the distal lobe of *C. reinhardtii* and human N-DRC is significantly thinner than that of *T. thermophila*, suggesting that the N-terminus of CCDC96 resides there (Fig. S5E and S6B). In addition, the WD40 domain of the N-terminus of CCDC96 is missing in the cryo-ET map of the *T. thermophila* CCDC96 deletion (Fig. S6B).

**Figure 3.**
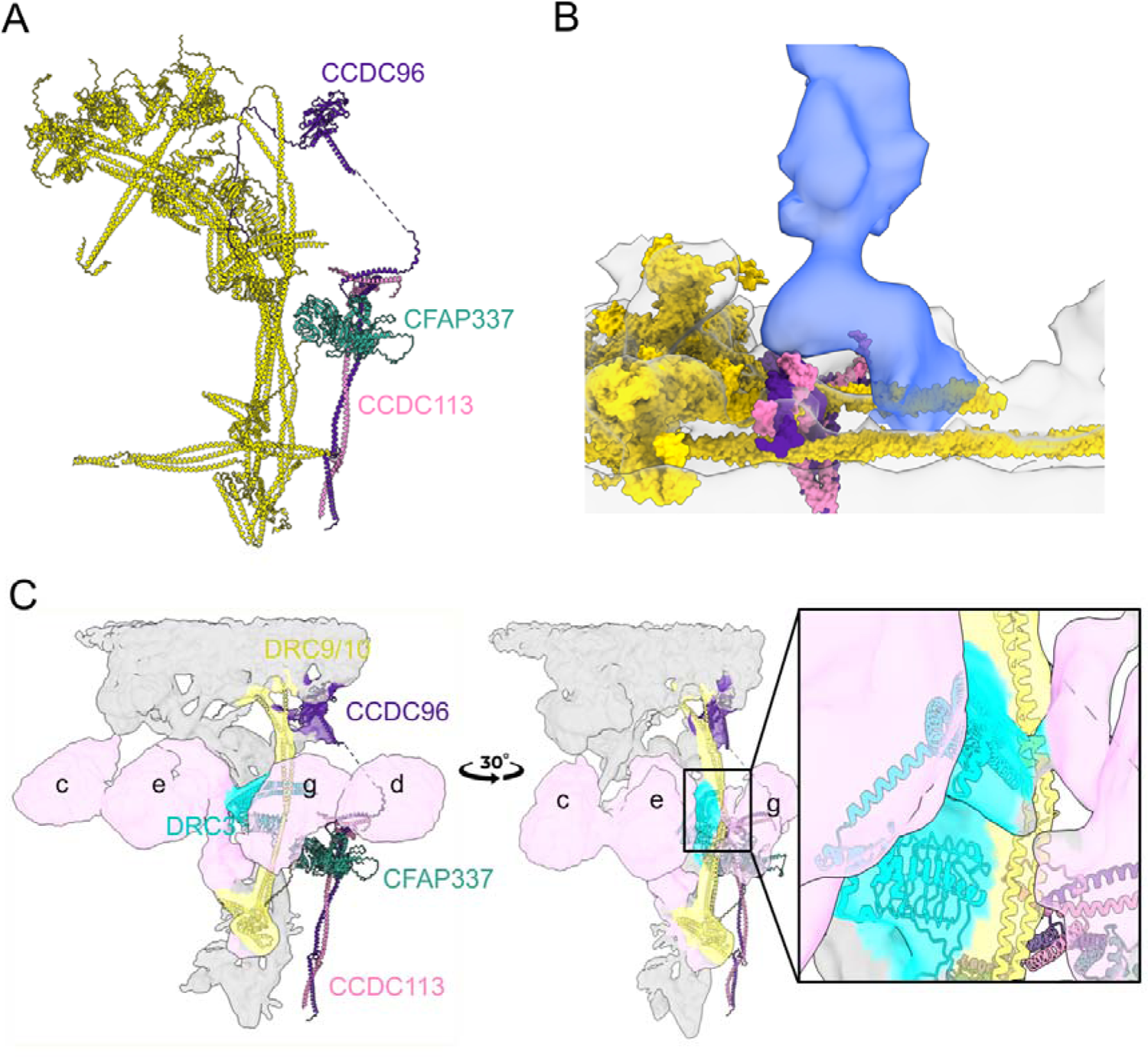
Structure of the CCDC96/113 complex. (A) The C-terminus of the CCDC96/113 coiled coil is located on the DMT, and the N-terminus is localized near the N-DRC linker domain. (B) The C-terminus of the CCDC96/113 complex interacts with the base of RS3. (The N-DRC and coiled-coil associated proteins except CCDC96/113 are represented in yellow). (C) The interactions of DRC3, DRC9/10 with IDAs (pink), and CCDC96/113 are demonstrated in the cryo-EM map and model.

The apparent proximity suggests that CCDC96 has crosstalk with the N-DRC. The N-termini of DRC9 and DRC3 interact with dynein e and g, respectively, suggesting that they are regulatory hubs for such dynein (Fig. 3C). Our subtomogram average of the 96-nm repeat indicates that the CCDC96/113 complex interacts with the base part of RS3 (Fig. 3B), as also suggested by a previous study [12]. Therefore, the CCDC96/113 complex may function as a signal transfer bridge for the N-DRC that runs from radial spoke RS3 to DRC10 and vice versa. Regulatory signals can be sent to dynein e through the N-terminus of DRC9. In addition, DRC10 may send signals to dynein g through DRC3. Therefore, the DRC9/10 coiled coil may be the most important regulatory hub in the N-DRC that regulates dyneins g and e by receiving the signal from CCDC96/113 (Fig. 3B-C).

We also found a crosslink from CFAP337A (UniProtID I7MM07, paralog CFAP337B UnitProtID I7MKT5) to the coiled-coil region of CCDC96. The AlphaFold2 predicted models of CFAP337A and CFAP337B contain two WD40 beta propellers and a bundled coiled-coil domain that fits unambiguously in a density close to the upper region of CCDC96 (Fig. S7A). Mass spectrometry analysis confirmed that both CFAP337A and CFAP337B are lost in CCDC96 knockout cells [12], suggesting that CFAP337A/B interacts with CCDC96. Our fitting of a predicted CFAP337 model, combined with cross-links and mass spectrometry data of the CCDC96 deletion mutant, demonstrated with good confidence that CFAP337 interacts directly with CCDC96. Our subtomogram average showed that the WD40 domain of CFAP337 mediates interaction with the stem of dynein d (Fig. S7B). CFAP337 may therefore be an IDA regulatory hub or linkage.

### The N-DRC regulates dynein activity through direct IDA interactions and a network of coiled-coil proteins that span the 96-nm repeat

The next step is to determine how the N-DRC regulates dynein activity and ciliary bending. To address this, we analyzed the single-particle and subtomogram averaged maps looking for any potential linkage from the N-DRC to the adjacent DMT and other axonemal complexes.

There are two observable connections from the N-DRC to the adjacent DMT, a clear connection from the distal lobe to protofilament B9 and a thin density from the middle of the linker part to protofilament B8 (Fig. 4A, black arrows). The rest of the N-DRC linker part does not show clear connections with the adjacent DMT and maintains a gap of ∼25 Å to protofilament B9. It is possible that the disordered regions of DRC2, DRC8, DRC9, DRC10, DRC11 and DRC12 can still contact the highly negative and flexible tails of tubulins from protofilament B10 of the neighboring DMT (Fig. 2D).

**Figure 4.**
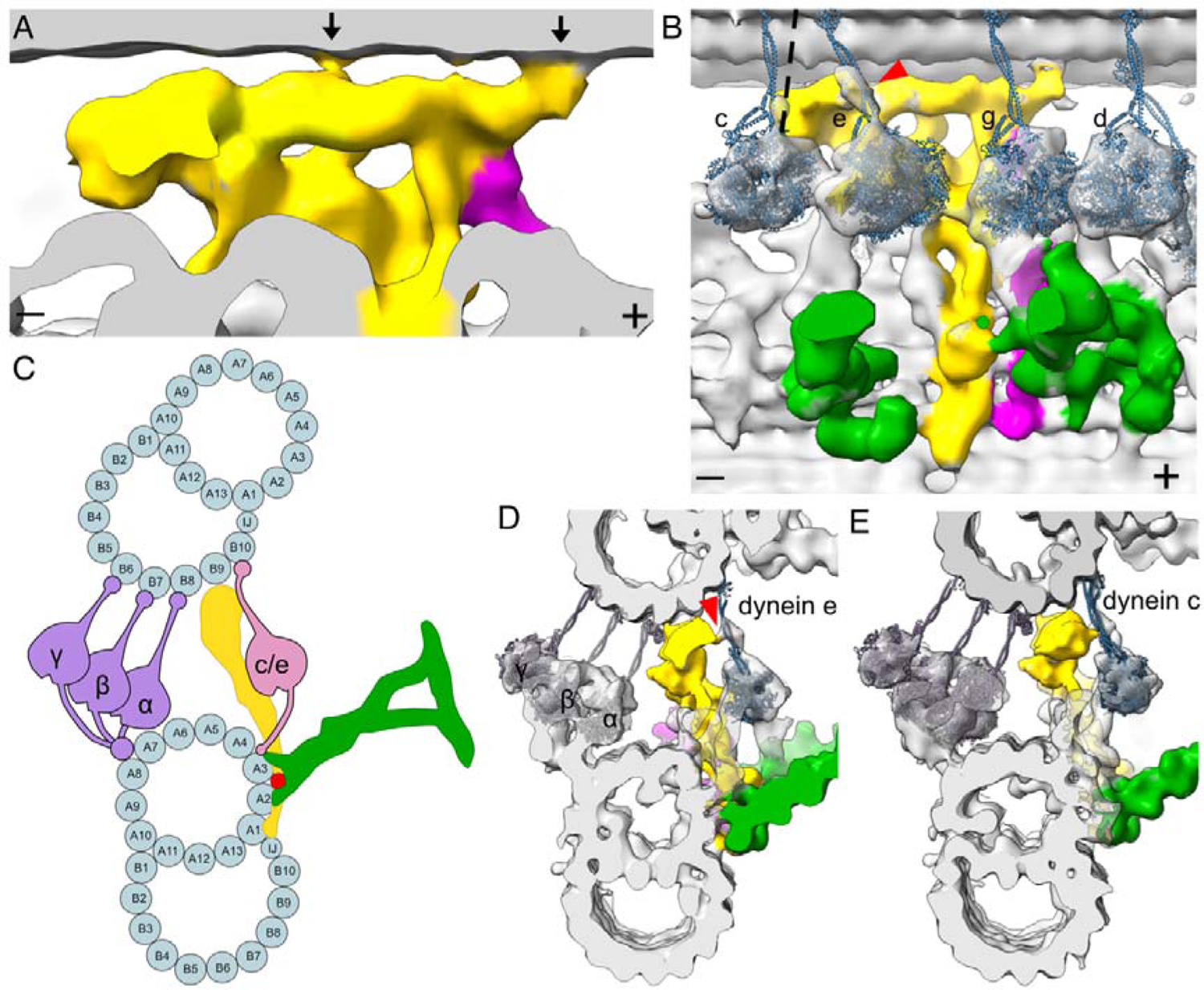
Interactions between the N-DRC and neighboring MTD. (A) Contacts of the N-DRC with the adjacent DMT in the distal lobe and middle part are indicated by black arrows. Signs (+) and (-) indicate the distal and proximal ends of the DMT. Colors: N-DRC: yellow; Radial spoke: green; CCDC96/113: dark purple. (B) Fitting of dynein heavy chain models into the IDA densities showing that the dynein e stalk interacts directly with the N-DRC proximal lobe (red arrowhead), while the dynein c stalk in the prepower stroke conformation (dotted line) might interact with the N-DRC proximal lobe. (C) A cross-sectional model of how dyneins and the N-DRC contact the neighboring DMT. (D) Cross-sectional view of the interaction between dynein e and the N-DRC. The same connection in (C) is indicated by the red arrowhead. (E) Cross-sectional view of the potential interaction between dynein c and the N-DRC.

We fitted atomic models of dyneins to the ODA and IDA densities (Fig. 4B, Fig. S7C, Methods). The stalks of ODA α, β, and γ dyneins contact protofilaments B5, B6 and B7, respectively, as reported previously [35] (Fig. 4C). Regarding IDA, the stalks of single-headed dyneins a, b, c, d, e, and g all bind to protofilament B10, while the stalks of double-headed inner arm dyneins fα and fβ bind to protofilament B9 (Fig. S7D, E). Interestingly, the stalk of dynein e appears to be in contact with the N-DRC proximal lobe corresponding to DRC8 and DRC11 (Fig. 4D). While the N-DRC and the stalk of dynein c do not seem to be in contact in the subtomogram average representing the postpower stroke state of dynein, the stalk of dynein c in the prepower stroke will move distally [36] (Fig. 4B and E) and therefore could be in contact with the N-DRC bulky proximal lobe. In addition to physical contact from DRC3 to dyneins e and g described in the previous section, we showed that the activity of the stalks of dyneins c and e can be regulated by the N-DRC linker region, particularly the DRC8/11 complex, through mechanical contact. Dynein c is a strong dynein and is particularly needed for cell movement at high viscosity [37] The swimming speed of the *C. reinhardtii* dynein c mutant (*ida9*) is only marginally reduced compared to wild-type cells in normal medium but greatly reduced in viscous media [37]. Similarly, the *C. reinhardtii* DRC11 mutant (*drc11*) swims only slightly slower than wild-type cells in normal medium [21]. Therefore, it might be possible that missing DRC11 affects dynein c regulation and has the same effect as missing dynein c.

Next, we looked for complexes in the same DMT that were in contact with the N-DRC. We identified a coiled-coil density located on the A3 and A4 protofilaments, which we named the A3A4 coiled coil (Fig. 5A). This density extends through the CCDC96/113 complex and interacts directly with the WD40 domain of CFAP337. In *C. reinhardtii*, knockout of CFAP57 leads to loss of CFAP337 [38], suggesting that CFAP57 is a strong candidate for the A3A4 coiled coil. There are four CFAP57 paralogs in *T. thermophila,* with CFAP57A and CFAP57C (UniProtID Q234G8 and W7WWA2) as the most abundant paralogs (Table. S2). The AlphaFold2 multimer predictions of CFAP57A and CFAP57C match well with the density in our map (Fig. S7F). CFAP57A was highly biotinylated in cells expressing CCDC96-HA-BirA* or CCDC113-HA-BirA* fusion proteins [12]. This further reinforces the likelihood that the A3A4 coiled coil is CFAP57. CFAP57 spans almost 96 nm along the DMT [38] with the N-terminal region connecting to the I1 intermediate chain/light chain domain and the C-terminus positioned proximal to the radial spoke RS1 in the next 96-nm repeat (Fig. 5B-D). Our subtomogram average and single-particle maps suggest that the C-terminal region of CFAP57 also interacts with RS3 (Fig. S7G) and that the CFAP57A/C dimer extends past CFAP337 through the N-DRC and toward the I1 intermediate chain/light chain complex. Our structural data demonstrated that the A3A4 coiled coil (CFAP57 heterodimer) mediates interaction with the base of dyneins g and d (Fig. S7H). In CFAP57-KO cells, dyneins g and d are decreased compared with those in WT cells [38, 39]. Therefore, CFAP57 can be an adaptor protein for the assembly of dynein g and d. Moreover, this protein can be a critical regulatory hub for dynein d and g since this coiled coil interacts directly with CCDC96/113, CFAP337, and the N-DRC.

**Figure 5.**
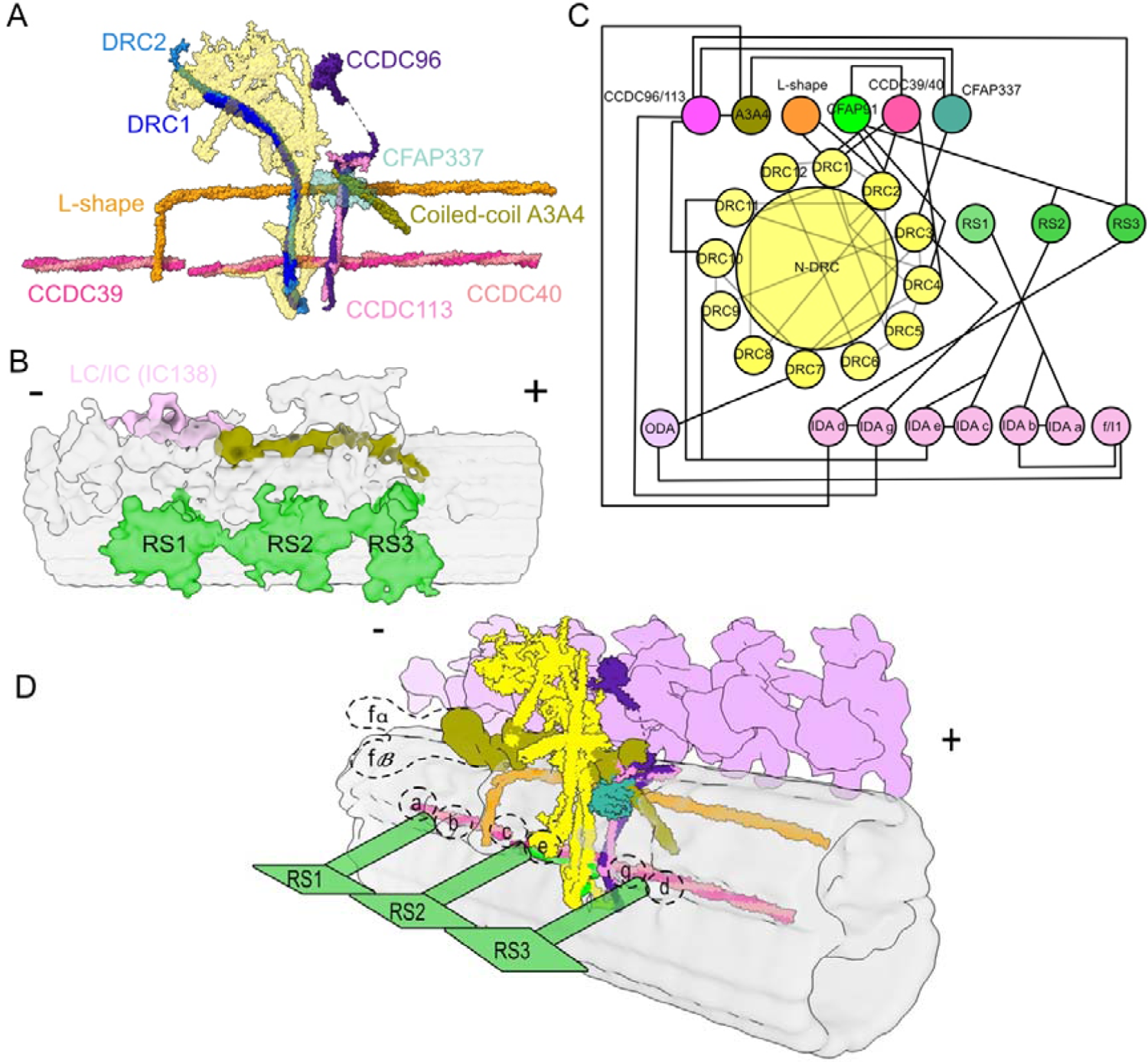
The coiled-coil network associated with the N-DRC. (A) N-DRC in complex with the coiled-coiled protein network. (B) The 96-nm repeat subtomogram averaged map of the DMT. (IDAs and ODAs are hidden for better visualization). The potential location of CFAP57. (C) Diagram of the N-DRC interactions with the coiled-coil network and IDAs and ODAs. The red line represents the N-DRC components. (D) Schematic view of the 96-nm repeating unit. The dashed shapes indicate the location of IDAs along the 96-nm repeat. Signs (+) and (-) indicate the distal and proximal ends of the DMT.

Our cryo-EM map also reveals the conservation of the L-shaped density discovered first in *C. reinhardtii* DMT [22]. The L-shaped coiled coil is located between RS1 and RS2 on protofilament A1 and extends to the junction of protofilaments A4 and A5. The L-shaped coiled coil interacts with CCDC39/40, DRC1/2, CCDC96/113 complexes, and CFAP337 (Fig. 5A, C, D). Overall, the N-DRC interacts with a network of coiled-coil proteins that can play a crucial role in the regulation of multiple dyneins.

Our subtomogram average does not show a linker density between the N-DRC and the ODA (Fig. S7B), as shown in *C. reinhardtii* [21]. In contrast to *T. thermophila*, DRC7 in *C. reinhardtii* has extra LRR and Kelch domains at its N-terminus. Therefore, the observed linker density can be the LRR and Kelch domains of CrDRC7. However, the flexible loop of TtDRC7 (N-terminal) might still interact with the IC/LC tower of ODA for N-DRC-ODA regulation.

### Mapping of pathogenic N-DRC human mutations allows the understanding of N-DRC assembly

In humans, mutations in N-DRC components such as scaffold proteins (DRC1, DRC2, and DRC4) disrupt cilia motility and lead to primary ciliary dyskinesia [40, 41]. To investigate the molecular basis of N-DRC-related ciliopathies, we mapped genetic mutations that cause ciliopathies on the human N-DRC model constructed based on our *T. thermophila* N-DRC model. DRC1 is an essential scaffold protein that binds to most DRC components. A Q118Term (DRC1) truncation mutation leads to a loss of the linker and base plate domain of DRC1, thereby destabilizing the entire N-DRC structure. The K686Term (DRC1) premature stop codon mutation truncates the binding region of DRC1 to CFAP20, which destabilizes the N-DRC base plate and causes severe defects in N-DRC assembly in primary ciliary dyskinesia patients (Fig. 6A). This suggests that linking DRC1 to CFAP20 is critical for maintaining the N-DRC structure on DMT.

**Figure 6.**
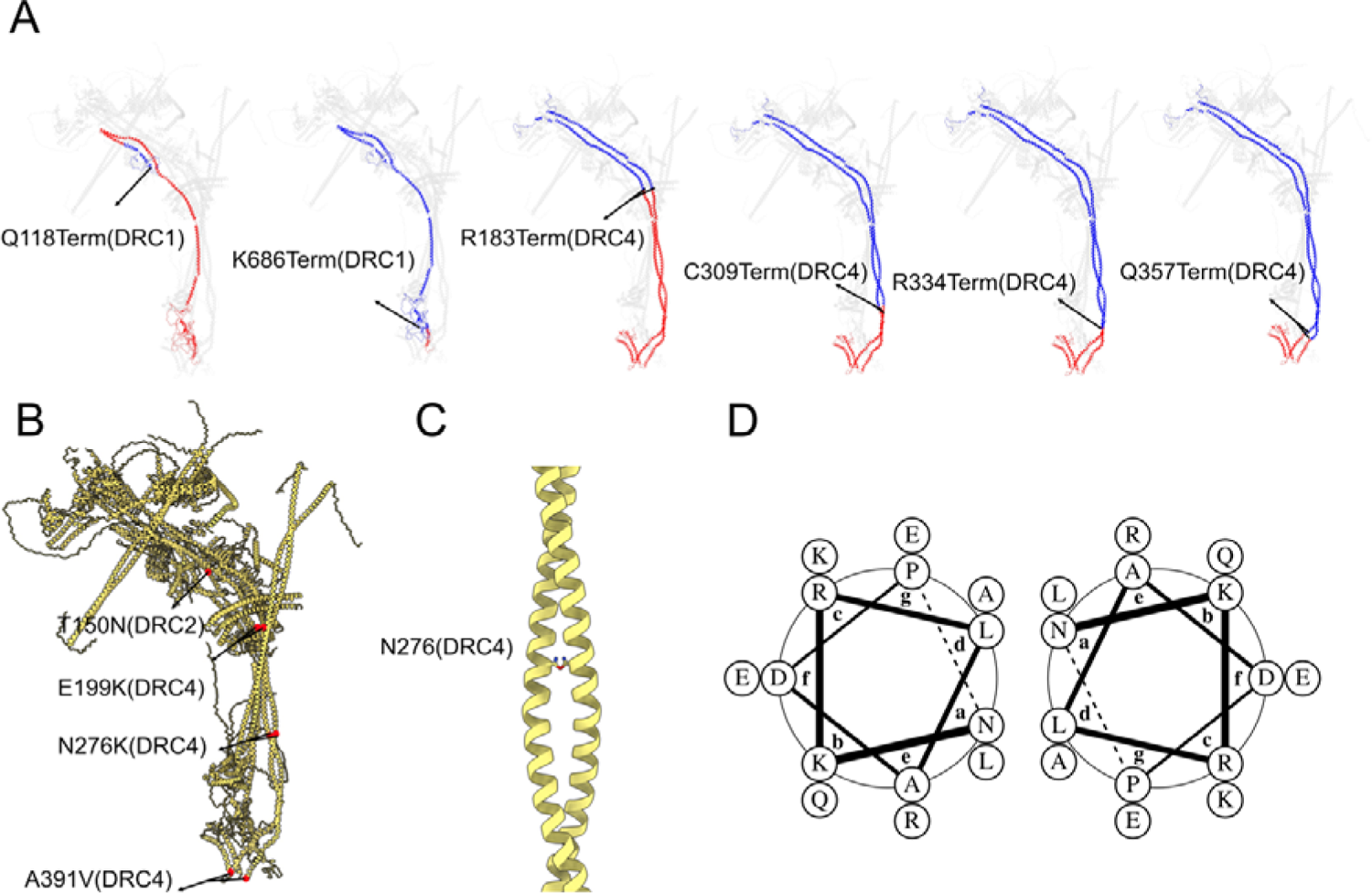
Genetic mapping of missense and truncated ciliopathies causing mutation in the human N-DRC model. (A) The pathogenic truncation mutations are shown on the human N-DRC model. The gray transparent proteins represent non-mutated proteins. The red color shows the truncated region, and the blue color depicts the remaining part of the protein. (B) Red circles indicate missense mutations in DRC4 and DRC2. Blue circles on DRC4 indicate benign missense mutations. (C) Interaction of side chain of asparagine residues in DRC4 coil coiled (D) Helical wheels of residue 276-289 segment of DRC4 coiled coil (sequence: NKRLADPLQKAREE).

Immunofluorescence analyses of respiratory cells from primary ciliary dyskinesia-affected individuals with truncation mutations in the DRC4 subunit (C309Term, R334Term, Q357Term) also revealed an N-DRC defect [42]. The premature stop codon mutations produce DRC4 variants lacking fragments at the N-DRC base plate, which in turn disrupts the interaction of DRC4 with ruler proteins (CCDC39/40), CFAP91, and tubulins, ultimately leading to a destabilization of the binding of the N-DRC to the DMT (Fig. 6A). The R183Term mutation in the DRC4 subunit also eliminates part of DRC4, which binds to DRC3 (Fig. 6A). As expected, this mutation led to delocalization of DRC3 subunits in the N-DRC [43].

It has been reported that missense mutations (N276K, A391V) in DRC4 cause disorganization of DMT and change the swimming velocity of cilia. Our model suggests that these mutations can lead to disorganization or disassembly of the DRC4 coiled-coil structure (Fig. 6B). A canonical coiled coil is composed of heptad repeats, labeled *abcdefg*, where hydrophobic amino acids at positions *a* and *d* are conserved [44]. Asparagine is among the most abundant polar inclusions at positions *a* or *d*, providing conformational dynamics at the coiled-coil interface [45]. Our structural data suggest that N276 is located at position *a* in the coiled-coil heptad (Fig. 6C-D). Therefore, mutation of N276 to a positively charged lysine can disrupt or deform the DRC4 coiled-coil structure.

In summary, scaffold proteins (DRC1, DRC2, DRC4) play a vital role in assembling the N-DRC since any defect in these subunits can cause severe ciliopathies. These proteins construct the N-DRC core structure and interact with most of the proteins in the base plate and linker regions. Due to the pivotal role of scaffold proteins in N-DRC assembly, any defect in these proteins can cause loss of the N-DRC structure and disrupt the contact of the N-DRC with the neighboring DMT. The contact of the N-DRC by electrostatic interactions with the neighboring DMT is essential for ciliary bending [34]. Previous studies in *C. reinhardtii* mutants of *DRC7* and *DRC11* show that the partially disrupted N-DRC still maintains contact with the neighboring DMT [21, 34]. Accordingly, if there are some contacts with the neighboring DMT, the bending motion will be affected but not completely disturbed. As a result, pathogenic N-DRC mutations are primarily found in scaffold proteins.

## Discussion

In this study, we revealed the molecular architecture of the N-DRC of *T. thermophila* by an integrative modeling approach. Importantly, our data identified an additional N-DRC component, DRC12. We localized CFAP337 and CFAP5 and demonstrated their interactions with the CCDC93/113 complex. The main scaffold of the N-DRC comprises coiled-coiled proteins, including DRC1/2 and DRC4A/4B complexes, which start from the base plate and span through the proximal lobe of the linker. These components play a pivotal role in stabilizing the N-DRC and contribute to mechano-regulatory signal transmission. The role of DRC1, DRC2 and DRC4 proteins as the core structural components of the N-DRC agrees with earlier analyses of the N-DRC in *C. reinhardtii* (N-C-term tagging SNAP, BCCP) [38] and biochemical studies [18].

In contrast to the radial spokes and ODA, which are preassembled in the cytoplasm and transported into the cilium [46–48], a recent study revealed that the N-DRC is likely assembled step-by-step in the cilia since DRC2 and DRC4 are transported and assembled independently in the cilia [49]. In accordance with our model and mass spectrometry analysis of different knockout DRC mutants [18, 24], scaffold proteins should assemble onto the DMT first before the other N-DRC subunits dock onto the core scaffold proteins. Loss of DRC3 in the *drc3* mutant strain led to the loss of part of the distal domain (based on our study, the distal domain that was lost in the *drc3* mutant represents DRC9/10 complex) [50]. We propose that DRC3 binds to DRC4 first, leading to the assembly of DRC9/10 heterodimers. As a result, DRC4 plays a crucial role in assembling the N-DRC distal domain.

Our model revealed that the N-DRC is connected to an extensive network of coiled-coil proteins on the DMT that span the 96-nm repeating unit. An essential aspect of coiled-coil protein function is mechanical stability against deformation forces [51]. Coiled-coil proteins can transmit mechanical loads such as the tails of myosin and kinesin motors or keratin bundles [51]. We hypothesized that the N-DRC conformation change during ciliary bending can be transmitted to the coiled-coil network by pulling force. To bend the axoneme, dyneins on one side should be inactive, while dyneins on the other side must be active [11]. During ciliary beating, this process should switch back and forth quickly, resulting in the high oscillating frequency of the cilium. When the cilium is in a straight conformation, the N-DRC is in its relaxed state (Fig. 7A). In this conformation, the N-DRC exerts no regulation on dynein arm activity on either side of the cilium. In the outward-facing curve of the cilium, the dyneins are in active states, leading to DMT sliding and N-DRC stretching due to the larger distance between neighboring DMTs (Fig. 7B). N-DRC stretching can pull on the highly elastic coiled-coil protein network. This can mechanically extend the length of coiled-coil proteins. As a result, the dynein arms can be regulated by the pulling of the coiled-coil proteins and switch to an inactive state. In active cilia, the N-DRC linker region is not well observed by cryo-ET, probably due to its flexibility [36]. As such, the extent of the deformation of the N-DRC linker region is still unclear and needs to be answered in the future.

**Figure 7.**
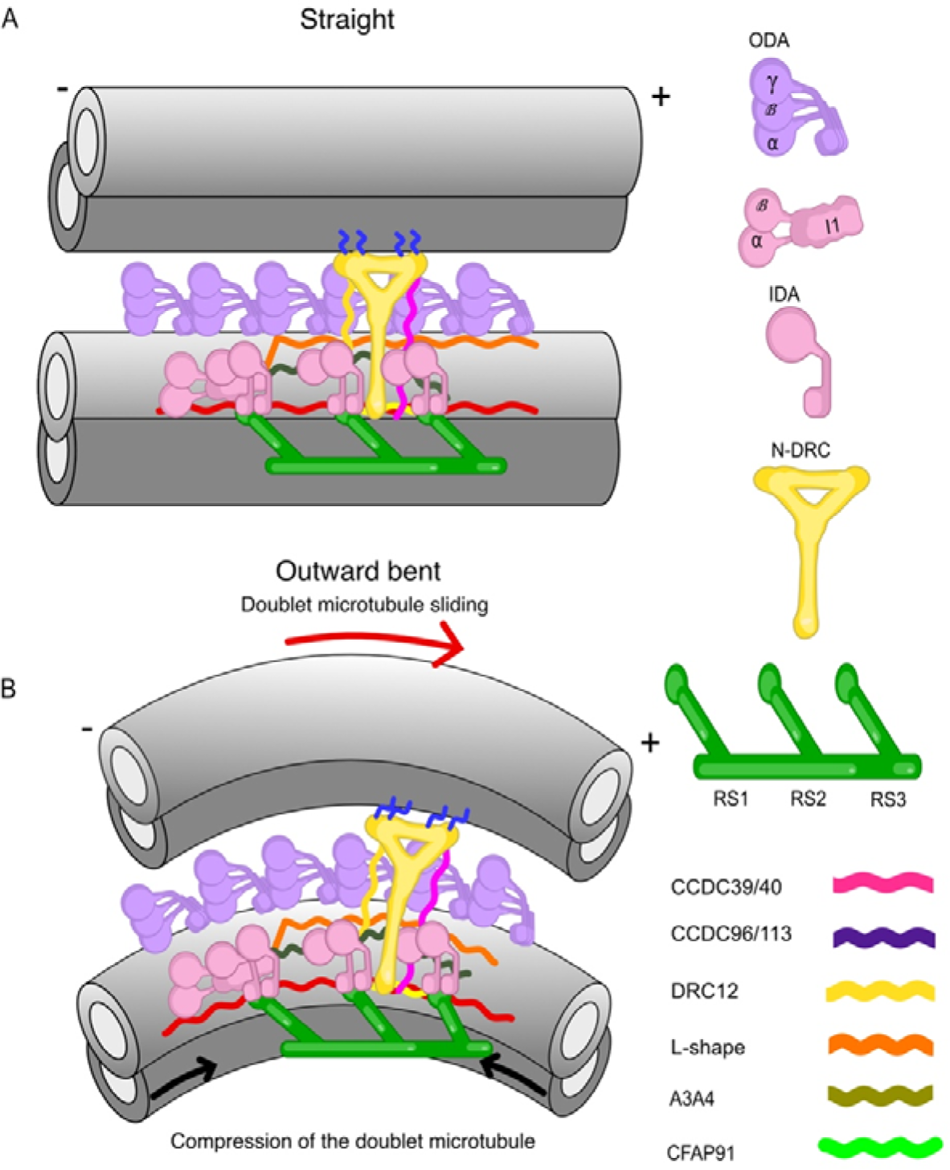
A model for the conformational change of the N-DRC and the coiled-coil network during ciliary bending. (A) Two neighboring DMTs are in their straight conformations. (B) The N-DRC pulls due to DMT sliding and the coiled coils on the surface of the DMT due to bending, which can lead to dynein activation/inactivation. Signs (+) and (-) indicate the distal and proximal ends of the DMT.

To summarize, our study sheds light on the molecular mechanism of dynein arm regulation through the N-DRC and other mechano-regulatory complexes. Our results allow us to better understand the molecular etiology of ciliopathies and the mechanism of ciliary dysfunction.

## Methods

*T. thermophila* growth, cilium isolation by dibucaine treatment, cryo-EM sample preparation, and cryo-EM data acquisition were performed similarly to our previous work [26, 27].

### T. thermophila culture

All *T. thermophila* strains in this study were grown in 1 L of SPP media in a shaker incubator at 30°C and 120 rpm until the cell density reached an OD of approximately 0.7 at 700 nm (Black, Dai et al. 2021).

### Cilia isolation

The *T. thermophila* culture was harvested by centrifugation at 700 × g for 10 min at room temperature. The cell pellet was resuspended in fresh SPP at room temperature to a total volume of 24 ml. One milliliter of SPP containing 25 mg of dibucaine was added to the cell solution for one minute for deciliation. Immediately, 75 ml of ice-cold SPP was added to the cell suspension. The solution was centrifuged at 2000 × g for 10 min at 4°C. The supernatant containing the cilia was then centrifuged at 17000 × g for 40 minutes at 4°C. The cilia-containing pellets were washed with ice-cold cilia wash buffer (50 mM HEPES at pH 7.4, 3 mM MgSO4, 0.1 mM EGTA, 1 mM DTT, 250 mM sucrose) and resuspended in 250 μl cilia wash buffer.

### Purification of DMTs

NP40 was added to a final concentration of 1.5% and incubated on ice for 45 minutes. After centrifugation for 10 minutes at 7800 g and 4°C, the intact axoneme with the membrane in the pellet was resuspended in cilia final buffer (50 mM HEPES at pH 7.4, 3 mM MgSO_4_, 0.1 mM EGTA, 1 mM DTT, 0.5% trehalose). To induce sliding disintegration for intact DMT purification, ADP was added to a final concentration of 0.3 mM and incubated at room temperature for 10 minutes. Then, ATP was added to a final concentration of 1 mM and incubated at room temperature for another 10 minutes to maximize sliding disintegration.

### Cryo-EM sample preparation

Chloroform was applied overnight to C-Flat Holey thick carbon grids for cleaning (Electron Microscopy Services #CFT312-100). For single-particle analysis, chloroform-treated and negatively glow-discharged (10 mA, 15 s) grids were placed inside the Vitrobot Mk IV (Thermo Fisher) chamber, and 4 μl of the DMT sample at 2.2 mg/mL was applied to the grid. The sample was incubated on the grid at 23°C for 15 seconds with 100% humidity, blotted with force 3 for 5 seconds, and then plunged frozen in liquid ethane.

### Cryo-EM data acquisition

Beam-shift single-particle cryo-EM data were collected by SerialEM [52] on a Titan Krios 300 keV FEG electron microscope (Thermo Fisher) equipped with a K3 Summit direct electron detector (Gatan, Inc.) and the BioQuantum energy filter (Gatan, Inc.) (Table 1). The beam shift pattern used to collect movies is four movies per hole and four holes per movement. The final pixel size is 1.370 Å/pixel. A total dose of 45 or 70 electrons per Å^2^ was radiated to each movie with over 40 frames. The defocus range was between −1.0 and −3.0 μm at an interval of 0.25 μm.

### Cryo-EM image processing of the 96-nm repeat unit of the DMT

The movies were motion-corrected and dose-weighted using MosionCor2 [53] in Relion 3.1.2 [54], and the contrast transfer function parameters were determined using GCTF [55]. The filaments were picked semi-automatically. Due to the importance of the top views of filaments, these views were picked manually using e2helixboxer [56]. The side views of the DMT were picked automatically using topaz [57] in Cryosparc 3.1 [58]. Four times binned particles of 128 × 128 pixels were extracted in Relion 3.1.2. The pre-alignment of 8-nm repeat particles was performed using the Iterative Helical Real Space Reconstruction script [59] using the SPIDER package [60]. In the next step, the particles were transferred to Frealign and refined [61].

After converting the alignment parameters to Relion 3.1.2 with an unbinned box size of 512 512 pixels, iterative per-particle-defocus refinement and Bayesian polishing of 8-nm repeat particles were performed. The signal of the tubulin lattice was subtracted to highlight the signal of MIPs for classification of 16-nm and 48-nm repeats. To obtain a 16-nm repeating unit, 3D classification was performed with two classes. The 16-nm repeat particles were subjected to 3D classification with three classes to obtain the 48-nm repeat. Refinement of the non-subtracted particles was performed to obtain a 48-nm structure. To obtain the 96-nm repeat of the DMT, K40R, MEC17, SB255, and CU428 data were merged since the N-DRC structure in the mutants is not different. Then, 3D classification with two classes with a mask focusing on the outside of the DMT was performed on the 48-nm repeat structure. The refinement of the 96-nm class containing N-DRC resulted in a global resolution of 3.6 to 4.0 Å.

To improve the local resolution of the base plate part, we performed local refinement using a mask that covered the base plate region and approximately two protofilaments attached to the base plate. Then, the map was enhanced by DeepEmhancer [62].

### Cryo-EM data processing of the N-DRC

A processing pipeline is shown in Figure S1. The N-DRC base plate was improved using focus refinement with one mask that included the base-plate part and CCDC96/113 complex on the DMT in Relion 3.1.2. To obtain the linker part, the particles of the 96-nm repeat were transferred to Cryosparc 3.1 to perform tubulin lattice signal subtraction, IDA subtraction, and centering of the N-DRC structure in the box using the volume alignment tool in Cryosparc 3.1. The star files were converted using the UCSF pyem package (https://github.com/asarnow/pyem) [63] to Relion 3.1.2. After consensus refinement of the entire N-DRC, the refined structure was subjected to 3D classification without alignment with five classes and a T value of 50. The best class with the highest number of particles and resolution was selected for further 3D refinement. Then, the N-DRC base plate was subtracted, and the box size was reduced to 150 pixels. The resulting particles were locally refined with three different masks covering different regions of the linker part. The final maps at 5.7 Å resolution were B-factor sharpened and enhanced by DeepEmhancer [62]. All maps and models were visualized using ChimeraX [64].

### Cryo-ET and subtomogram averaging

The data for cryo-electron tomography and the 96-nm repeat map (EMD-29667) of the wild-type cilium come from a previous study [27]. To improve the resolution of the N-DRC, local refinement was performed using a mask that covers the N-DRC structure, resulting in a 21 Å resolved map.

### Identification of N-DRC proteins in *T. thermophila*

To identify the DRC subunits of *T. thermophila*, the *C. reinhardtii* DRC subunits (obtained from previous studies and the UniProt and Phytozome databases [25, 65]) were blasted against the local database of the *T. thermophila* axoneme proteome, which was built using proteins identified in mass spectrometry data from the *T. thermophila* axoneme [27, 66, 67]. Additionally, the exponentially modified protein abundance index (emPAI) was considered another important factor in identifying DRC components (Table S2). DRC4, DRC6 and DRC11 all have two paralogs, A and B. In the case of DRC6 and DRC11, in which one paralog is significantly more abundant, we used the more abundant paralogs DRC6A and DRC11A for modeling (Table S1-2).

The intra-molecular and inter-molecular cross-links from the N-DRC components came from *in situ* cross-linking data of *T. thermophila* cilia [28] (Table S3 and S4).

### AlphaFold2 Structure Prediction

Using the AlphaFold2 Google Colab notebook [68, 69], we predicted all N-DRC components individually. To localize proteins and model heterodimer and homodimer proteins, we utilized AlphaFold2 Multimer [70] implemented in the Cosmic 2 computer cluster [71] and Google Colab notebook [72]. All models were predicted using the full sequence. The model was truncated and refined to fit into the cryo-EM map.

### Localization and modeling of N-DRC proteins

For base plate modeling, we used two different methods, including multiple sequence alignment and AlphaFold2 prediction. For the N-DRC base plate, multiple sequence alignment of DRC1, DRC2, CFAP91, and CFAP20 was performed against *C. reinhardtii* homologs. In the next step, the PDB structure of C. reinhardtii (PDB 7JU4) was modified into the sequence of *T. thermophila*. For DRC4A/4B, and CCDC96/113, we used AlphaFold2 prediction. The models were then fitted into the map using the UCSF ChimeraX function fitmap [64]. Next, the models were modeled using Coot [68] and real space refined in Phenix [69].

To localize DRC3, DRC9, DRC10, and DRC7, we performed AlphaFold2 Multimer protein complex prediction, which confirmed that DRC9 and DRC10 tightly bind together and form a heterodimer that interacts with DRC3 (Fig. S2F). The structure of DRC3 was resolved as a bundle of coiled-coil proteins connected to a bundle of beta sheets with a loop in the middle of the linker domain. This allowed us to fit the high-confidence AlphaFold2 structure into the density (Fig. S2C). The DRC5 density was resolved as a leucine-rich density that interacts with the linker domain of DRC1/2, consistent with a previous study [21]. The high-confidence AlphaFold2 multimer model of DRC1 and DRC5 confirms the interaction between these proteins (Fig. S2C, F). In our cryo-EM map, the resolved density of the DRC9/10 coiled coil has a distinct N-terminal fold, consistent with the molecular model predicted by the AlphaFold2 multimer. Therefore, we were able to fit the N-terminal regions and coiled-coil regions of the DRC9/10 coiled coil entirely within our map (Fig. S2E). The sufficient resolution of the linker domain also allowed us to fit the AlphaFold2 multimer model of DRC1/2 and DRC4A/4B complexes into our map.

For the identification and localization of other proteins, only proteins with at least two kinds of localization evidence are discussed in the work. The evidence includes cross-links to well-localized proteins, biotinylated in BioID of known proteins, highly confident AlphaFold2 multimer prediction with known proteins, presence in a pull-down assay with known bait proteins, missing in knockout mutants of known proteins and good fitting in low-resolution maps.

The cross-linking data indicated that DRC7 is cross-linked with DRC10 and DRC3. Hence, to localize DRC7, we performed AlphaFold2 multimer prediction of DRC7, DRC10, and DRC3 (Fig. S2F). In addition, the linker part of DRC1/2 was also localized using AlphaFold2 Multimer (Fig. S2A).

To determine the location of DRC8 and DRC11 on the N-DRC structure, we utilized Assembline as an integrative modeling software, AlphaFold2 Multimer, Hdock protein docking server, and Crosslink MS/MS data. A high-confidence AlphaFold2 Multimer prediction and the high score of the Hdock docking server and Assembline [73–77] enabled us to fit the DRC8/11 complex in the N-DRC proximal lobe (Fig. S2F). Using the UCSF ChimeraX function fitmap and Assembline Global optimization function, two copies of DRC8 and one copy of DRC11 localized in our map as a subcomplex. The fit libraries were created with 100000 searches in the cryo-EM map using the fitmap tool of the UCSF Chimera [78] with the requirement of at least 60% of the input structure being covered by the EM map envelope defined at a low-density threshold. This resulted in 500-2800 alternative fitting in the cryo-EM map. Then, the resulting fit libraries of the DRC8 and DRC11 structures were used as input for the simultaneous fitting of DRC8 and DRC11 using the cryo-EM map. The fitting was performed using simulated annealing Monte Carlo optimization that generates alternative configurations of the fits precalculated as above.

DRC4 in *T. thermophila* has two paralogs compared with *C. reinhardtii*. Our cryo-EM map shows two coiled-coil densities for the DRC4 paralogs, in which one coil has a longer loop DRC4A (Q23YW7) with a particular fold and the other coil has a shorter loop DRC4B (I7LT80) (Fig. S5C). To identify how these two paralogs interact with each other and to distinguish which density on our map belongs to these proteins, we performed AlphaFold2 multimer prediction. Interestingly, AlphaFold2 Multimer predicted the dimer with high accuracy, allowing us to localize the proteins (Fig. S5C).

### Visualization of axonemal dyneins and DMT interaction

To visualize the dynein and axonemal dynein interaction, the isolated *T. thermophila* outer dynein arm model (PDB 7k58) was fitted into the 96-nm repeat subtomogram average (EMD-29667). For IDA, the head and stalk domains of *T. thermophila* dynein heavy chain DYH10 (UniProtID I7LTP7) were modeled using the Phyre2 server [79]. This model was then used to fit into all eight different inner arm dynein positions (a, b, c, d, e, g, fα and fβ). The fitting is based on the head and orientation of the density of the stalks in the subtomogram average.

### Genome modifications

To engineer *Tetrahymena* cells expressing C-terminally −3HA or –HA-BirA*-tagged DRC fusion proteins under the control of the respective native promoter, the FAP44 open reading frame and 3’UTR fragments were removed from FAP44-3HA and FAP44-HA-BirA* plasmids [80] using MluI and BamHI or PstI and XhoI sites, respectively, and replaced by approximately 1 kb fragments of the C-terminal part of the open reading frame without a stop codon and 3’UTR amplified using wild-type genomic DNA as a template and Phusion Hot Start II high-fidelity DNA polymerase (Thermo Fisher Scientific Baltics, Lithuania) and primers listed in Table S5.

To engineer *Tetrahymena* cells overexpressing C-terminally GFP- or HA-tagged proteins under the control of a cadmium-inducible MTT1 promoter, the entire open reading frame without the stop codon was amplified with the addition of the MluI and BamHI restriction sites at the 5’ and 3’ ends, respectively, using Phusion Hot Start II high-fidelity DNA polymerase (Thermo Fisher Scientific Baltics, Lithuania) and the primers listed in Table S5. The PCR fragments were cloned using MluI and BamHI restriction enzymes into modified versions of the MTT1-GFP or MTT1-HA plasmids [81], enabling the integration of the transgene into the *BTU1* locus and selection of transformed *Tetrahymena* cells based on paromomycin resistance (neo2 cassette) [82]. *Tetrahymena* cells were transformed, and transgenes were assorted as described in detail [12, 30].

### BioID assay

For the BioID assay, cells were grown to a density of 2 × 10^5^ cells/ml, starved for 14-18 hours in 10 mM Tris–HCl buffer, pH 7.5, and incubated in the same buffer supplied with 50 μM biotin for 4 h at 30°C. Next, the cells were spun down and deciliated, and the collected cilia were resuspended in 0.5 ml of axoneme stabilization buffer (20 mM potassium acetate, 5 mM MgSO_4_, 20 mM HEPES, pH 7.5, 0.5 mM EDTA with protease inhibitors (Complete Ultra EDTA-free; Roche, Indianapolis, IN)). After incubation of cilia for 5 min on ice in the same buffer supplied with 0.2% NP-40, the axonemes were pelleted by centrifugation at 21,100 *× g* for 10 min at 4°C and lysed for 1 hour (0.4% SDS, 50 mM Tris–HCl, pH 7.4, 500 mM NaCl, 1 mM DTT with protease inhibitors) at RT. After centrifugation (8000 *× g* at 4°C), the supernatant was collected and diluted with three volumes of 50 mM Tris–HCl buffer, pH 7.4. Collected proteins were incubated overnight with 100 μl of streptavidin-coupled Dynabeads (Dynabeads M-280 Streptavidin, Thermo Fisher Scientific, Waltham, MA, USA) at 4°C. After washing with buffer containing 15 mM Tris–HCl, pH 7.4, 150 mM NaCl, 0.1% SDS, and 0.3 mM DTT at 4°C, the biotinylated, resin-bound proteins were analyzed by mass spectrometry (Laboratory of Mass Spectrometry, Institute of Biochemistry and Biophysics, PAS, Warsaw, Poland) and by Western blotting using Pierce High Sensitivity-streptavidin-HRP (Thermo Scientific, Rockford, IL) diluted 1:40,000 in 3% BSA/TBST.

### Pull-down analyses

For the pull-down assay, cells carrying a transgene enabling overexpression of *Tetrahymena* proteins were grown overnight to the mid-log phase in SPP medium and diluted to a density of 2-2.5 × 10^5^ cells/ml, and overexpression was induced by the addition of cadmium chloride to a final concentration of 2.5 µg/mL. After 2–3 h, cells were collected, washed in 10 mM Tris-HCl buffer, pH 7.5, and solubilized in two-fold concentrated modified RIPA buffer (50 mM Tris-HCl, pH 7.5, 300 mM NaCl, 2% NP-40, 2% sodium deoxycholate, 10% glycerol with protease inhibitors).

After spinning down at 100,000 × g for 30 min at 4°C, the supernatant obtained from cells expressing GFP-tagged proteins was diluted 1:4 with dilution buffer (25 mM Tris-HCl, pH 7.5, 150 mM NaCl, 5% glycerol with protease inhibitors), and approximately 250 µg of protein was incubated with anti-GFP-conjugated magnetic agarose beads (ChromoTek GFP-Trap^®^ Magnetic Agarose) for 1-1.5 hours. Next, beads were washed (5 × 5 min) with Wash buffer (25 mM Tris-HCl, pH 7.5, 150 mM NaCl, 0.2% NP-40.5% glycerol with protease inhibitors) and incubated with approximately 250 µg of supernatant containing HA-tagged proteins prepared as described above for GFP fusion proteins. After 1-1.5 hours of incubation at room temperature, followed by washing with wash buffer, the bead-bound proteins were analyzed by Western blotting using anti-GFP (1:60,000, Abcam) and anti-HA antibodies (1:2000).

## Acknowledgments

We thank Drs. Kelly Sears, Mike Strauss, Kaustuv Basu (Facility for Electron Microscopy Research at McGill University) for helping with data collection, Dr. Jan Kosinski for helping to set up Assembline and Drs. Maureen Wirschell and Muneyoshi Ichikawa for critically reading the manuscript. KHB is supported by grants from the Canadian Institutes of Health Research (PJT-156354) and Natural Sciences and Engineering Research Council of Canada (RGPIN-2022-04774). EMM acknowledges support from the Welch Foundation (F1515) and U.S. National Institutes of Health (R01 HD085901). DW is supported by a grant from the National Science Centre, Poland (OPUS21 2021/41/B/NZ3/03612).

## Author contributions

Conceptualization: KHB, DW. Methodology: AG, SM, CLM, BN, CSB, SKY, TL, OP, MJ, MVP. Formal analysis: AG, SM, CLM. Investigation: AG, SM, CLM, BN, CSB, SKY, TL, OP, MJ, MVP, EMM. Resources: DW, KHB, EMM. Writing–original draft: KHB, DW, AG. Writing–review & editing: KHB, DW, AG, EMM.

## Data availability

The data generated in this study are available in the following databases: PDB-XXXX (*Tetrahymena* DRC model)

EMD-XXXXX (combined 96-nm *Tetrahymena* doublet)

EMD-XXXXX (combined N-DRC base plate part *Tetrahymena*) EMD-XXXXX (combined N-DRC linker part *Tetrahymena*)

EMD-XXXXX (focused refinement of the DRC from the *Tetrahymena* WT subtomo)

