## Supplementary Information for "Integrated modeling of the Nexin-dynein regulatory complex reveals its regulatory mechanism"

### Supplementary Figures

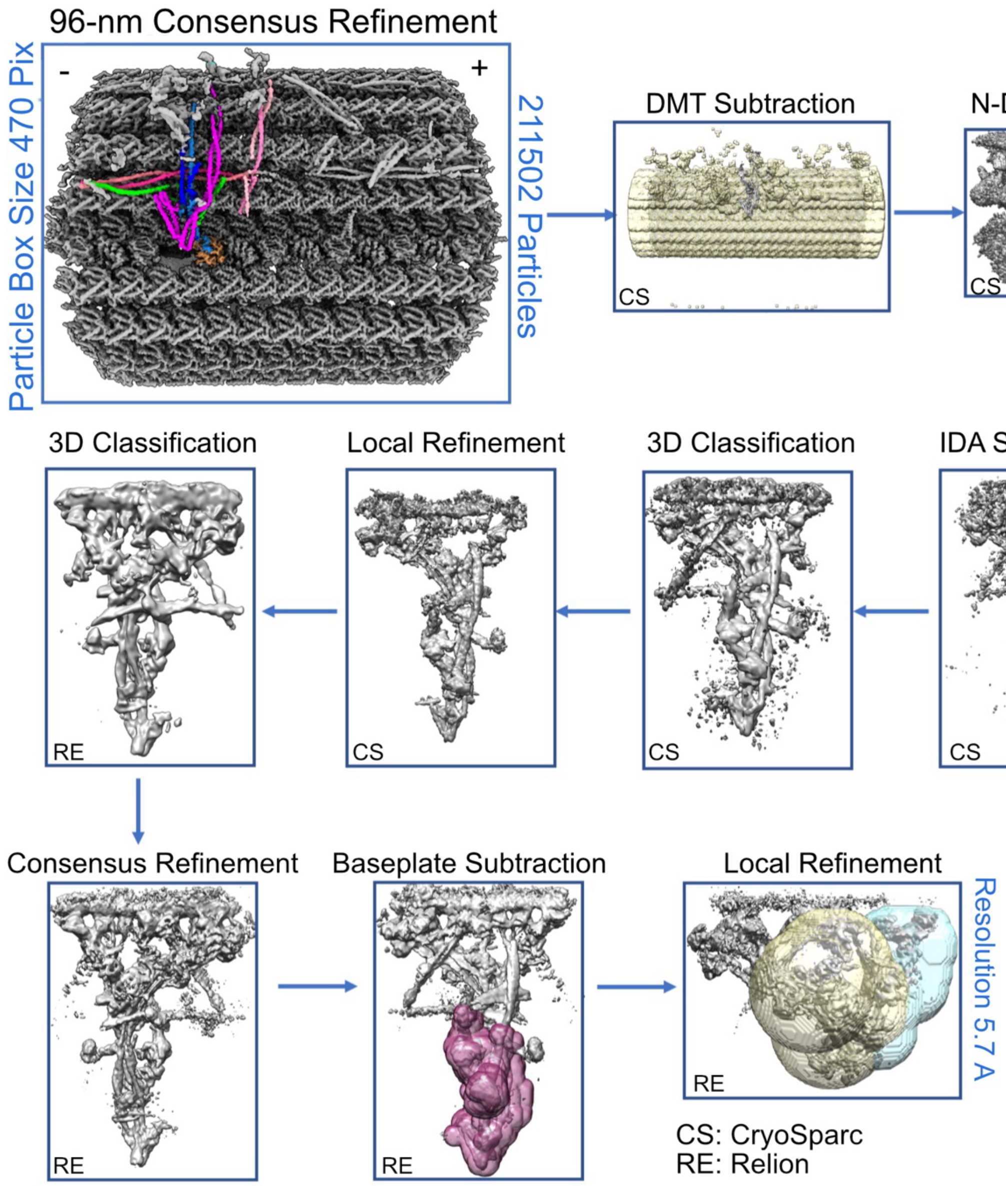


**Figure S1. The workflow of reconstruction of base plate and linker regions of N-DRC.**

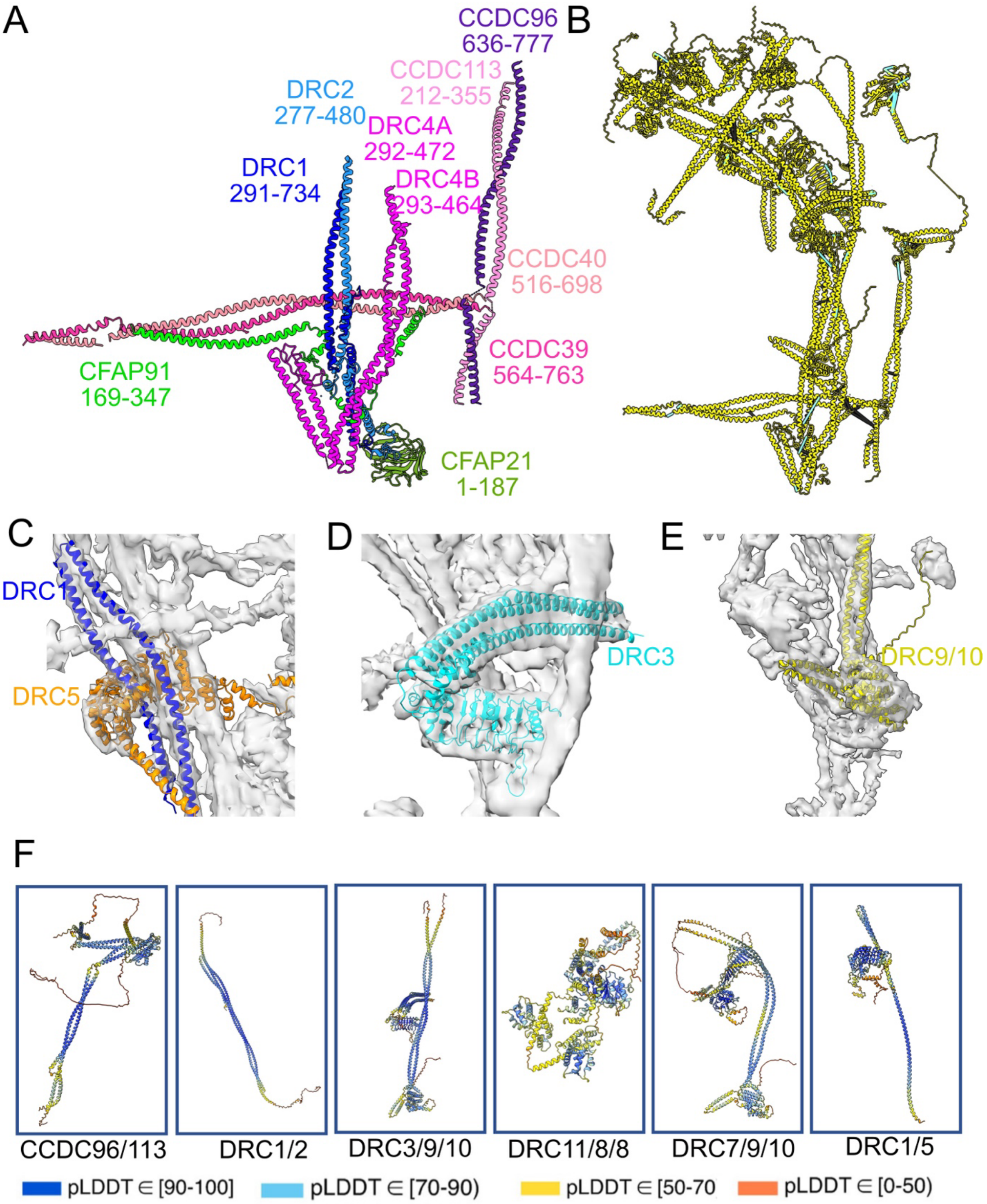


**Figure S2. AlphaFold2 modelling of all N-DRC components.** (A) The sequence range of the baseplate part is modelled using Coot. (B) The intra-molecular (cyan bars) and inter-molecular (black bars) cross-links are identified in the N-DRC. (C) DRC3 is fitted as a bundle of coiled-coil structures in the N-DRC cryo-EM map. In our N-DRC cryo-EM map, DRC1 and DRC5 are fitted. (D) In our N-DRC cryo-EM map, DRC1 and DRC5 are fitted. (E) The DRC9/10 N-terminus is fitted to the N-DRC Cryo-EM map. (F) The AlphaFold2 Multimer models of the N-DRC and CCDC96/113 components are coloured based on the pLDDT score where blue indicates the highest score, and orange represents the lowest score.

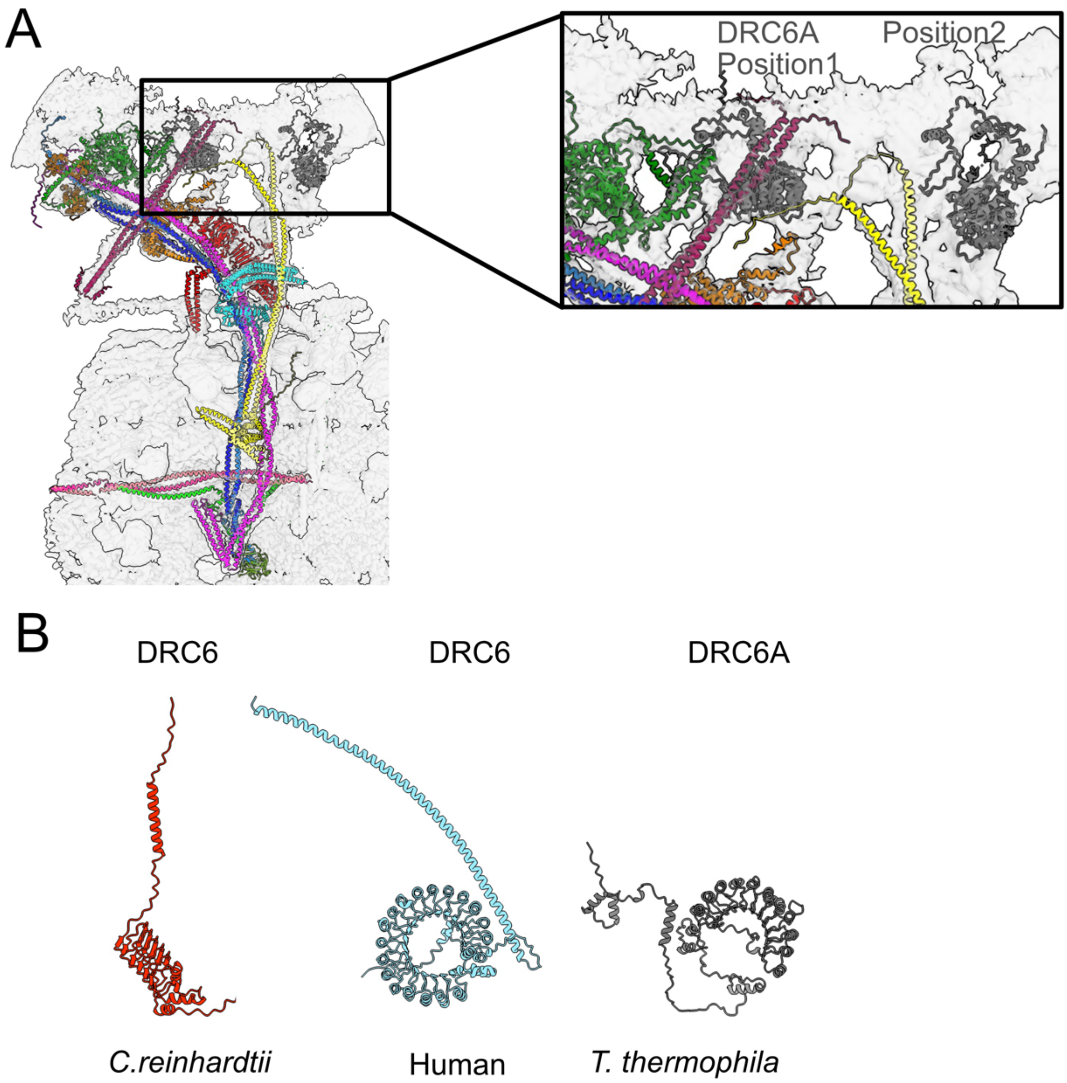


**Figure S3. Localizing DRC6A in the N-DRC cryo-EM map.** (A) Two potential locations for DRC6A in the cryo-EM map. (B) The structure of DRC6 in three different species.

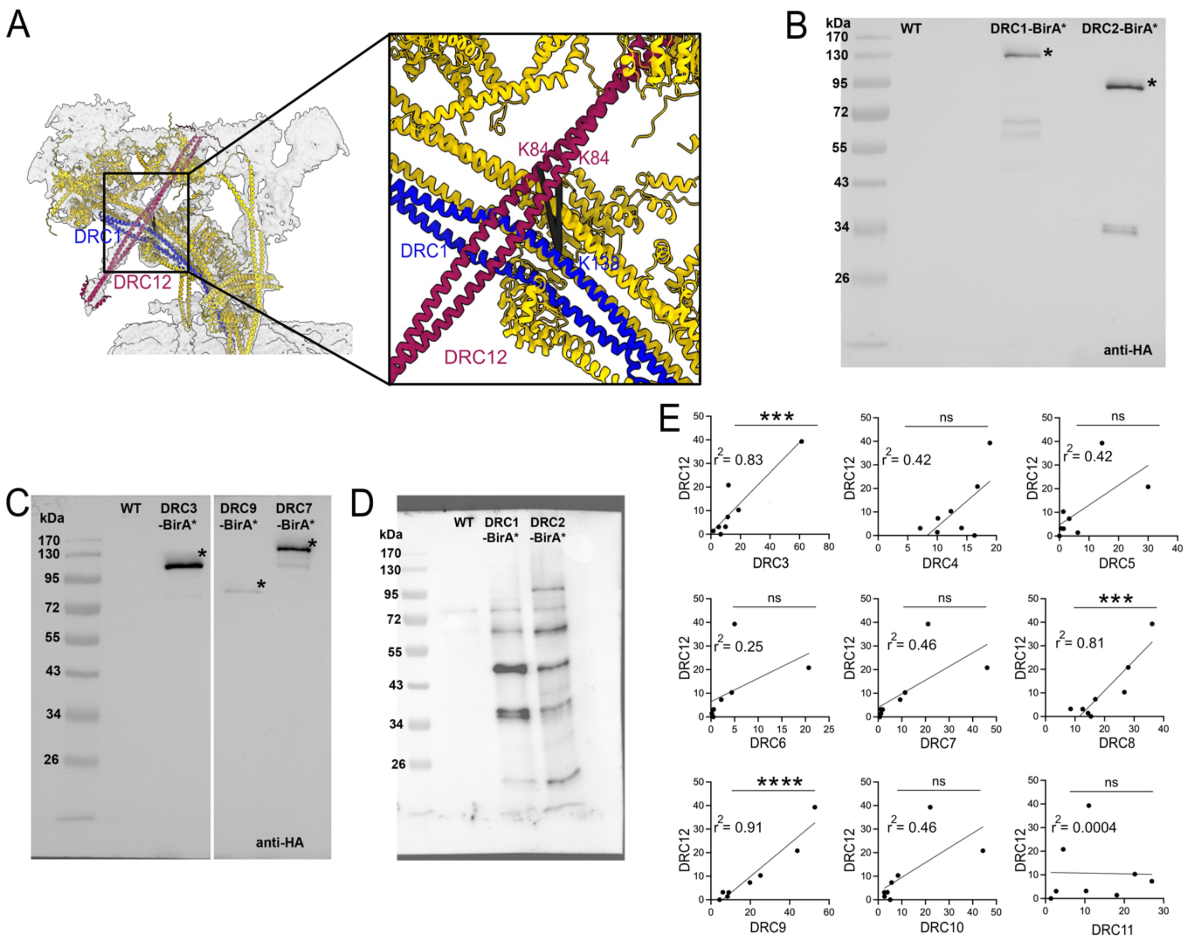


**Figure S4. CCDC153 is a DRC protein.** (A) Localization of DRC12 using cross-link MS/MS. The length of the cross-link between DRC1 and DRC12 is 27.26 and 30.61 Å. (B-C) Western blots with HA antibody of purified cilia from *WT*, *DRC1-HA-BirA** and *DRC2-HA-BirA** cells (B) and *DRC3-HA-BirA*, DRC7-HA-BirA** and *DRC9-HA-BirA** (C) indicate the baits are in the cilia. (D) Western blots using Streptavidin-HRP of purified cilia from *WT, DRC1-HA-BirA** and *DRC2-HA-BirA** cells after 4 h incubation with 50 μM biotin at 30°C showed that biotinylation of ciliary components happens. (E) Correlation graphs of consensus normalized RNA expression levels for DRC components vs DRC12. The coefficient of determination (R2) of the linear regression is indicated. *$p < 0.05$; ***$p < 0.001$;****$p < 0.0001$; ns: Not Significant.

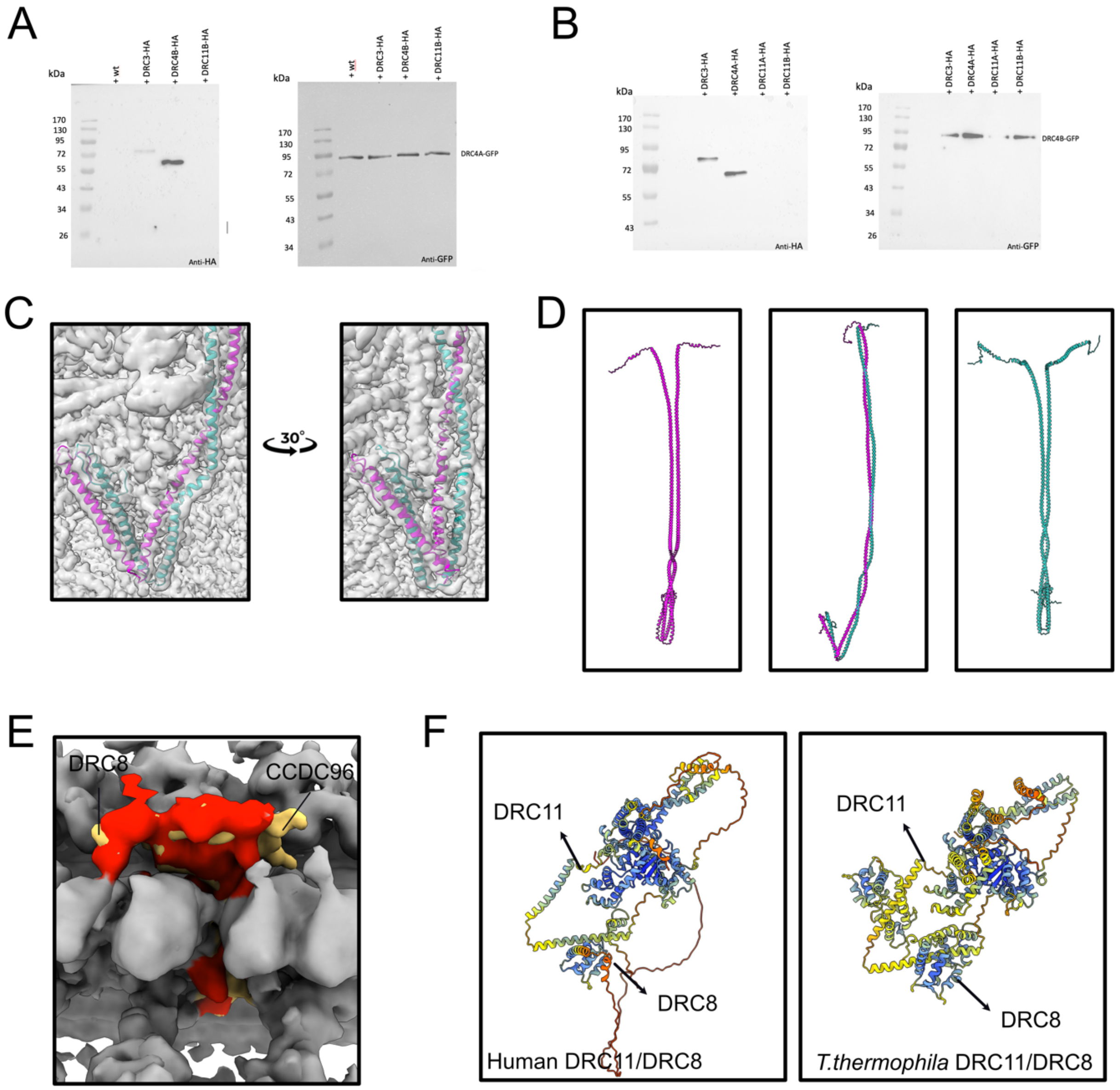


**Figure S5. The structure of DRC4 of the N-DRC.** (A, B) Pull down showing proteins pulled down by (A) DRC4A-GFP or (B) DRC4B-GFP bound to anti-GFP-beads. (C) The green and magenta colours indicate DRC4A and DRC4B, respectively. DRC4B contains a longer loop compared to DRC4A. D) The AlphaFold2 Multimer model of DRC4B/DRC4B, DRC4A/DRC4B, and DRC4A/DRC4A. The baseplate region of DRC4A/B shows that DRC4B has a longer loop than another protein. (E) Superimposed map of human (red) and *T. thermophila* (yellow). The larger density in the proximal lobe and distal lobe of the *T. thermophila* map indicates the second copy of DRC8 and N-terminus of CCDC96, respectively, which the human map lacks this density. (F) The AlphaFold2 Multimer prediction of DRC11/DRC8 subcomplex for humans and *T. thermophila*. The AlphaFold2 prediction indicates only one copy of DRC8 binds to DRC11 in humans. The AlphaFold2 models are coloured based on the pLDDT score where blue represents the highest confidence and orange represents the lowest confidence.

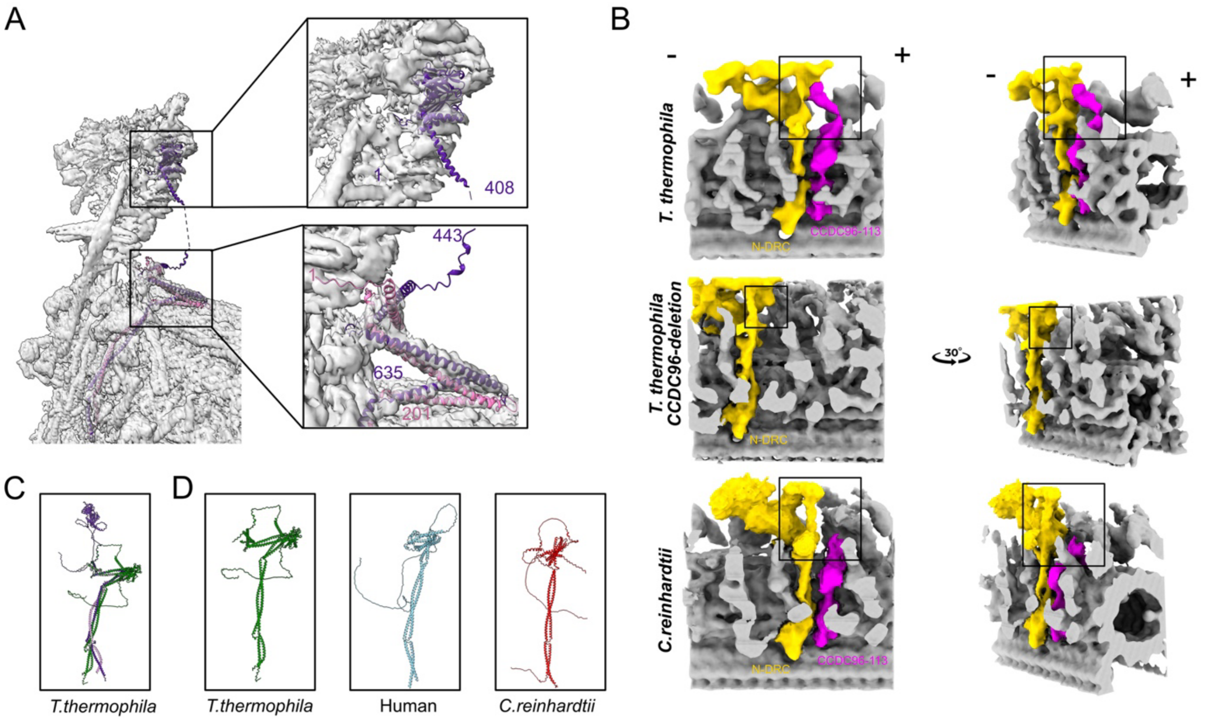


**Figure S6. The structure of CCDC96-113 in three different species.** (A) Fitting the N-terminal domain of CCDC96 in the map. The numbers represent number of amino acids on each protein. (B) The cryo-ET map of *T. thermophila* wild-type, *T. thermophila* CCDC96-KO, and *C. reinhardtii.* The inner dynein arms are deleted for better visualization. Signs (+) and (-) indicate the distal and proximal ends of the DMT. (C) Green colour indicates AphaFold2 multimer structure before truncation and refinement. The purple colour demonstrates protein after truncation and refinement. (D) CCDC96 has an extra globular domain compared to *C. reinhardtii* and Human.

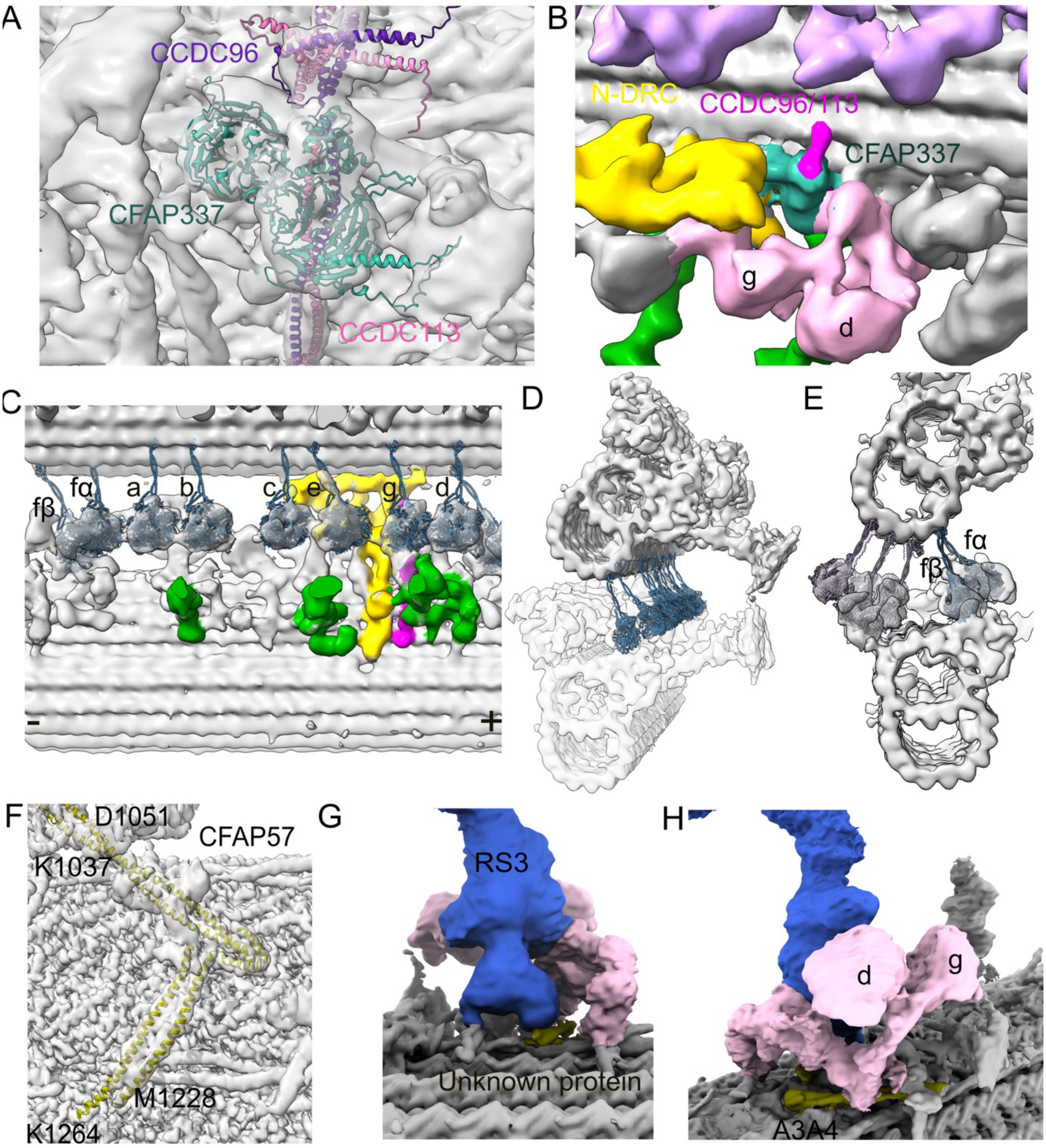


**Figure S7. The subtomogram averaging map of WT *T. thermophila*.** (A) Fitting the CFAP337 in cryo-EM map. (B) The map shows that CFAP337 interacts with CCDC96/113 and stems of dyneins g and d. (C) Fitting of dynein heavy chains into the IDA densities in the subtomogram average. Colors: N-DRC: yellow; RS: green; CCDC96/113: purple. Signs (+) and (-) indicate the distal and proximal ends of the DMT. (D) The stalks of inner arm dyneins contact protofilament B10 and B9. (E) Cross-sectional view shows that the stalks of dyneins fα and fβ binding to protofilament B9. (F) The map indicates A3A4 coiled coil interact with stem of dyneins d and g. (G) The fitting of AlphaFold2 predicted model of CFAP57A/C (UniprotID Q234G8 and W7WWA2) in our cryo-EM map. (H) The unknown protein interacts with the base of the RS3.

### Supplementary Tables

**Table S1.** The method of localizing and modelling of DRC components in cryo-EM map.

| DRC subunits | Uniprot ID | Gene name | AA-region Coot Modelling | AA-region AlphaFold2 localizing |
|---|---|---|---|---|
| DRC1 | Q229S1 | TTHERM_01345750 | 291-734 | 292-826 |
| DRC2 | Q24DJ0 | TTHERM_00971830 | 277-480 | 278-576 |
| DRC3 | I7MG46 | TTHERM_00316370 | — | 31-575 |
| DRC4A | Q23YW7 | TTHERM_00857910 | 293-464 | 1-294 |
| DRC4B | I7LT80 | TTHERM_00649240 | 292-472 | 1-293 |
| DRC5 | Q24C31 | TTHERM_00697450 | — | 1-461 |
| DRC6A | I7M1R2 | TTHERM_00339720 | — | 188-531 |
| DRC7 | I7MLZ4 | TTHERM_00473320 | — | 1-852 |
| DRC8 | W7WX86 | TTHERM_001232262 | — | 1-185 |
| DRC9 | Q23S05 | TTHERM_00625950 | — | 1-372 |
| DRC10 | A4VD15 | TTHERM_00535929 | — | 1-434 |
| DRC11A | I7LWE3 | TTHERM_00151820 | — | 1-862 |
| DRC12 | Q22RH5 | TTHERM_00016120 | — | 1-187 |
| CCDC96 | I7M6D6 | TTHERM_00529650 | 636-777 | 104-408, 442-626 |
| CCDC113 | Q22KK0 | TTHERM_00312810 | 212-355 | 1-202 |
| CCDC39 | Q23BW0 | TTHERM_00227220 | 533-763 | 101-308, 530-310 |
| CCDC40 | Q233L0 | TTHERM_00391400 | 516-698 | 53-257, 259-480 |
| CFAP91 | I7LWP7 | TTHERM_00578560 | 169-347 | — |
| CFAP20 | Q22NU3 | TTHERM_00418580 | 1-187 | — |
| CFAP337A | I7MM07 | TTHERM_00475060 | — | 36-865 |
| CFAP57A | Q234G8 | TTHERM_00105300 | — | 1005-1047 |
| CFAP57C | W7WWA2 | TTHERM_00052490 | — | 993-1037 |

**Table S2.** The homologs of N-DRC subunits of *T. thermophila* and human based on N-DRC subunits of *C. reinhardtii*.

| **DRC subunits** | ***C. reinhardtii* Uniprot ID** | **Human Uniprot ID** | ***T. thermophila* Uniprot ID** | **Resistant to salt extract** | **emPAI score** |
|---|---|---|---|---|---|
| DRC1 | P0DL09 | Q96MC2 | Q229S1 | Yes | 1.457 |
| DRC2 | A8JB22 | Q8IXS2 | Q24DJ0 | Yes | 3.253 |
| DRC3 | A8IVX2 | Q9H069 | I7MG46 | Yes | 3.685 |
| DRC4A | Q7XJ96 | O95995 | Q23YW7 | Yes | 2.499 |
| DRC4B | Q7XJ96 | O95995 | I7LT80 | Yes | 3.567 |
| DRC5 | A8HMZ4 | Q5JU00 | Q24C31 | Yes | 1.521 |
| DRC6A | A8JHD7 | Q8NEE6 | I7M1R2 | Yes | 1.188 |
| DRC6B | A8JHD7 | Q8NEE6 | I7MIQ9 | No | 0.202 |
| DRC7 | A0A2K3CXC4 | Q8IY82 | I7MLZ4 | Yes | 2.389 |
| DRC8 | A8J3A0 | Q5VUJ9 | W7WX86 | Yes | 1.823 |
| DRC9 | A0A2K3E6B1 | Q9H095 | Q23S05 | Yes | 2.442 |
| DRC10 | A8J0N6 | Q96DY2 | A4VD15 | Yes | 1.883 |
| DRC11A | A0A2K3DJI6 | Q86XH1 | I7LWE3 | Yes | 2.326 |
| DRC11B | A0A2K3DJI6 | Q86XH1 | W7XI81 | No | 0.316 |
| DRC12 | A0A2K3DW17 | Q494R4 | Q22RH5 | Yes | 2.401 |
| CFAP337A | A0A2K3DIJ7 | A0A804HJG6 | I7MM07 | No | 1.694 |
| CFAP337B | A0A2K3DIJ7 | A0A804HJG6 | I7MKT5 | No | 3.399 |
| CCDC96 | A0A2K3DWC0 | Q2M329 | I7M6D6 | Yes | 1.697 |
| CCDC113 | A0A2K3CQ22 | Q9H0I3 | Q22KK0 | Yes | 2.933 |
| CCDC39 | A0A0A1H1F6 | Q9UFE4 | Q23BW0 | No | 3.230 |
| CCDC40 | A0A0A1GYE8 | Q4G0X9 | Q233L0 | No | 20.723 |
| CFAP91 | A0A2K3DJU2 | Q7Z4T9 | I7LWP7 | No | 0.319 |
| CFAP20 | A8IU92 | Q9Y6A4 | Q22NU3 | No | 0.648 |
| CFAP57A | A0A2K3DTF5 | A0A087WVY5 | Q234G8 | No | 1.225 |
| CFAP57C | A0A2K3DV21 | A0A087WVY5 | W7WWA2 | No | 0.924 |
| CFAP57B | A0A2K3DRH4 | A0A087WVY5 | Q23CU1 | No | 0.609 |
| CFAP57D | | A0A087WVY5 | Q22N63 | No | 0.402 |

**Table S3**. Intermolecular cross link data identified from N-DRC.

| Protein 1 | Uniprot ID | Residue number | Protein 2 | Uniprot ID | Residue number |
|---|---|---|---|---|---|
| CCDC39 | Q23BW0 | 660 | CFAP91 | I7LWP7 | 218 |
| CCDC39 | Q23BW0 | 613 | DRC2 | Q24DJ0 | 342 |
| CFAP91 | I7LWP7 | 343 | CCDC96 | I7M6D6 | 750 |
| CFAP91 | I7LWP7 | 347 | CCDC96 | I7M6D6 | 750 |
| CFAP91 | I7LWP7 | 343 | CCDC40 | Q233L0 | 542 |
| CFAP91 | I7LWP7 | 343 | CCDC40 | Q233L0 | 531 |
| DRC4A | Q23YW7 | 123 | DRC1 | Q229S1 | 150 |
| CCDC153 | Q22RH5 | 83 | DRC1 | Q229S1 | 138 |
| DRC1 | Q229S1 | 178 | DRC4A | Q23YW7 | 149 |
| DRC1 | Q229S1 | 180 | DRC2 | Q24DJ0 | 172 |
| DRC1 | Q229S1 | 258 | DRC2 | Q24DJ0 | 247 |
| CCDC96 | I7M6D6 | 665 | CCDC113 | Q22KK0 | 245 |
| CCDC96 | I7M6D6 | 697 | CCDC113 | Q22KK0 | 274 |
| CCDC96 | I7M6D6 | 679 | CCDC113 | Q22KK0 | 259 |
| CCDC113 | Q22KK0 | 346 | CCDC96 | I7M6D6 | 761 |
| CCDC113 | Q22KK0 | 108 | CCDC96 | I7M6D6 | 532 |
| CCDC113 | Q22KK0 | 52 | CCDC96 | I7M6D6 | 532 |
| CCDC113 | Q22KK0 | 152 | CCDC96 | I7M6D6 | 573 |
| DRC4B | I7LT80 | 332 | CCDC40 | Q233L0 | 542 |
| DRC4B | I7LT80 | 322 | DRC4A | Q23YW7 | 315 |
| DRC4B | I7LT80 | 83 | DRC4A | Q23YW7 | 73 |
| DRC4B | I7LT80 | 201 | DRC4A | Q23YW7 | 194 |
| DRC4B | I7LT80 | 55 | DRC11A | I7LWE3 | 551 |
| DRC4B | I7LT80 | 56 | DRC11A | I7LWE3 | 562 |
| DRC4B | I7LT80 | 378 | DRC2 | Q24DJ0 | 367 |
| DRC4B | I7LT80 | 261 | DRC2 | Q24DJ0 | 262 |
| DRC4B | I7LT80 | 218 | DRC2 | Q24DJ0 | 255 |
| DRC10 | A4VD15 | 346 | DRC7 | I7MLZ4 | 250 |
| DRC3 | I7MG46 | 377 | DRC7 | I7MLZ4 | 566 |
| DRC3 | I7MG46 | 355 | DRC7 | I7MLZ4 | 221 |
| CFAP91 | I7LWP7 | 343 | DRC4B | I7LT80 | 332 |
| DRC2 | Q24DJ0 | 228 | DRC4B | I7LT80 | 233 |

**Table S4.** Intramolecular cross-links identified from N-DRC.

| Protein 1 | Uniprot ID | Residue Number | Protein 2 | Uniprot ID | Residue number |
|---|---|---|---|---|---|
| DRC4A | Q23YW7 | 100 | DRC4A | Q23YW7 | 93 |
| DRC5 | Q24C31 | 397 | DRC5 | Q24C31 | 387 |
| DRC5 | Q24C31 | 91 | DRC5 | Q24C31 | 83 |
| DRC1 | Q229S1 | 187 | DRC1 | Q229S1 | 180 |
| CCDC96 | I7M6D6 | 351 | CCDC96 | I7M6D6 | 289 |
| CCDC96 | I7M6D6 | 368 | CCDC96 | I7M6D6 | 255 |
| CCDC96 | I7M6D6 | 380 | CCDC96 | I7M6D6 | 375 |
| CCDC96 | I7M6D6 | 351 | CCDC96 | I7M6D6 | 289 |
| CCDC96 | I7M6D6 | 255 | CCDC96 | I7M6D6 | 375 |
| CCDC96 | I7M6D6 | 255 | CCDC96 | I7M6D6 | 380 |
| CCDC96 | I7M6D6 | 263 | CCDC96 | I7M6D6 | 339 |
| CCDC96 | I7M6D6 | 380 | CCDC96 | I7M6D6 | 269 |
| CCDC96 | I7M6D6 | 255 | CCDC96 | I7M6D6 | 375 |
| CCDC96 | I7M6D6 | 263 | CCDC96 | I7M6D6 | 339 |
| CCDC96 | I7M6D6 | 351 | CCDC96 | I7M6D6 | 289 |
| CCDC113 | Q22KK0 | 108 | CCDC113 | Q22KK0 | 100 |
| CCDC113 | Q22KK0 | 52 | CCDC113 | Q22KK0 | 108 |
| CCDC113 | Q22KK0 | 280 | CCDC113 | Q22KK0 | 284 |
| DRC11A | I7LWE3 | 309 | DRC11A | I7LWE3 | 313 |
| DRC11A | I7LWE3 | 559 | DRC11A | I7LWE3 | 562 |
| DRC11A | I7LWE3 | 831 | DRC11A | I7LWE3 | 562 |
| DRC7 | I7MLZ4 | 488 | DRC7 | I7MLZ4 | 464 |
| DRC7 | I7MLZ4 | 433 | DRC7 | I7MLZ4 | 464 |
| DRC7 | I7MLZ4 | 576 | DRC7 | I7MLZ4 | 681 |
| DRC7 | I7MLZ4 | 576 | DRC7 | I7MLZ4 | 566 |
| DRC10 | A4VD15 | 277 | DRC10 | A4VD15 | 285 |
| DRC10 | A4VD15 | 239 | DRC10 | A4VD15 | 246 |
| DRC3 | I7MG46 | 386 | DRC3 | I7MG46 | 392 |
| DRC3 | I7MG46 | 463 | DRC3 | I7MG46 | 302 |
| DRC3 | I7MG46 | 53 | DRC3 | I7MG46 | 88 |
| DRC3 | I7MG46 | 119 | DRC3 | I7MG46 | 27 |
| DRC3 | I7MG46 | 385 | DRC3 | I7MG46 | 392 |

**Table S5.** Mass spectrometry-based identification of ciliary proteins biotinylated in cells expressing the C-terminally -HA-BirA* tagged DRC proteins. Numbers X/Y: (X) number of all identified peptides (in a Mascot program, all significant matches), (Y) number of all unique peptide sequences (in a Mascot program, significant sequences).

| | | Exp 1 | | | Exp 2 | | | | Exp 3 | | |
|---|---|---|---|---|---|---|---|---|---|---|---|
| | | WT | DRC1-HA-BirA* | DRC2-HA-BirA* | 0 | DRC1-HA-BirA* | DRC2-HA-BirA* | DRC3-HA-BirA* | WT | DRC7-HA-BirA* | DRC9-HA-BirA* |
| DRC1 | TTHERM_01345750 | 1/1 | 31/20 | 16/14 | 0 | 9/7 | 10/7 | 0 | 0 | 0 | 0 |
| DRC2 | TTHERM_00971830 | 0 | 17/10 | 12/10 | 0 | 11/7 | 26/15 | 2/2 | 0 | 0 | 0 |
| DRC3 | TTHERM_00316370 | 0 | 18/10 | 17/11 | 0 | 3/2 | 6/4 | 17/11 | 0 | 0 | 0 |
| DRC4A | TTHERM_00857910 | 2/2 | 23/14 | 20/11 | 0 | 10/9 | 21/13 | 2/2 | 0 | 1/1 | 0 |
| DRC4B | TTHERM_00649240 | 3/1 | 23/12 | 23/13 | 0 | 5/4 | 10/8 | 2/2 | 0 | 1/1 | 2/1 |
| DRC5 | TTHERM_00697450 | 0 | 0 | 1/1 | 0 | 0 | 1/1 | 0 | 0 | 2/2 | 0 |
| DRC6A | TTHERM_00339720 | 0 | 0 | 0 | 0 | 0 | 1/1 | 0 | 0 | 0 | 0 |
| DRC6B | TTHERM_00522350 | 0 | 0 | 0 | 0 | 0 | 0 | 0 | 0 | 0 | 0 |
| DRC7 | TTHERM_00473320 | 0 | 19/11 | 16/14 | 0 | 3/2 | 3/3 | 1/1 | 0 | 43/25 | 0 |
| DRC8 | TTHERM_001232262 | 0 | 0 | 0 | 0 | 1/1 | 0 | 0 | 0 | 0 | 0 |
| DRC9 | TTHERM_00625950 | 6/4 | 12/10 | 16/8 | 0 | 1/1 | 7/6 | 3/3 | 0 | 0 | 7/4 |
| DRC10 | TTHERM_00535929 | 0 | 0 | 0 | 0 | 6/4 | 13/9 | 3/3 | 0 | 0 | 0 |
| DRC11A | TTHERM_00151820 | 2/2 | 13/8 | 14/12 | 0 | 0 | 0 | 0 | 0 | 0 | 0 |
| DRC11B | TTHERM_000268319 | 0 | 0 | 0 | 0 | 0 | 1/1 | 2/2 | 1/1 | 1/1 | 0 |
| DRC12 | TTHERM_00016120 | 0 | 0 | 0 | 0 | 3/2 | 1/1 | 0 | 0 | 0 | 0 |
| CCDC96 | TTHERM_00529650 | 1/1 | 29/18 | 8/6 | 0 | 1/1 | 0 | 0 | 0 | 0 | 0 |
| CCDC113 | TTHERM_00312810 | 4/4 | 27/16 | 17/11 | 0 | 45/20 | 1/1 | 1/1 | 0 | 0 | 0 |
| CFAP337A | TTHERM_00475060 | 0 | 3/2 | 1/1 | 0 | 0 | 0 | 0 | 0 | 0 | 0 |
| CFAP337B | TTHERM_00218510 | 0 | 1/1 | 0 | 0 | 0 | 0 | 2/2 | 0 | 0 | 0 |
| CFAP91 | TTHERM_00578560 | 0 | 4/3 | 6/4 | 0 | 0 | 0 | 0 | 0 | 0 | 0 |
| CFAP57A | TTHERM_00105300 | 0 | 45/26 | 24/18 | 0 | 29/21 | 7/6 | 0 | 0 | 0 | 0 |
| CFAP57B | TTHERM_00052490 | 0 | 4/3 | 2/2 | 0 | 0 | 0 | 0 | 0 | 0 | 0 |
| CFAP57C | TTHERM_00214710 | 0 | 0 | 0 | 0 | 0 | 0 | 0 | 0 | 0 | 0 |
| CFAP57D | TTHERM_000681920 | 0 | 0 | 0 | 0 | 32/23 | 13/9 | 0 | 0 | 0 | 0 |
| CCDC39 | TTHERM_00227220 | 0 | 2/2 | 2/2 | 0 | 0 | 0 | 0 | 0 | 0 | 0 |
| CCDC40 | TTHERM_00391400 | 0 | 3/2 | 0 | | 0 | 0 | 0 | 0 | 0 | 0 |
| CFAP20 | TTHERM_00418580 | 11/7 | 25/7 | 25/8 | | 1/1 | 2/1 | 0 | 0 | 0 | 0 |

**Table S6.** Primers used in this study.

| Gene | Primer's name | Primers' nucleotide sequence |
|---|---|---|
| expression of C-terminally –HA-BirA* tagged DRC proteins under the control of respective native promoter | | |
| *DRC1* | DRC1-Bir-cod-MluI-F | AAAT**ACGCGT**GAGGAAGCAAGAAAATAACTTCTTTT G |
| | DRC1-Bir-cod-BamHI-R | AATT**GGATCC**TCTTTGTCCTTAATCTATATTAATGAGTCTTG |
| | DRC1-Bir-3UTR-PstI-F | AAAT**CTGCAG**CCTACTCTAAAAAAATTATTCTCCTGATAAAA |
| | DRC1-Bir-3UTR-XhoI-R | AATT**CTCGAG**AATCTTCAAAAAGTAAGTAGATTGATAGATAG |
| *DRC2* | DRC2-Bir-cod-MluI-F | AAAT**ACGCGT**CATGCCCATAATAAGAATAGAAAGTAGGA |
| | DRC2-Bir-cod-BamHI-R | AATT**GGATCC**CTAGAATGGAGGAGCTCTTTATCC |
| | DRC2-Bir-3UTR-PstI-F | AAAT**CTGCAG**CACTCTATCCATTAATTACATTTGTATCTG |
| | DRC2-Bir-3UTR-XhoI-R | AATT**CTCGAG**CGAACTTTTGGTTGAGTTCAAAAG |
| *DRC3* | DRC3-Bir-cod-MluI-F | AATT**ACGCGT**GGGAGAAAGAAGCTGCTAATG |
| | DRC3-Bir-cod-BamHI-R | AATT**GGATCC**ATTTTCATGATCGGATTCATCATCTC |
| | DRC3-Bir-3UTR-PstI-F | AATT**CTGCAG**AGATCTATCTATCTATTGACTTAATTGATTTGT |
| | DRC3-Bir-3UTR-XhoI-R | AATT**CTCGAG**ATCGATCTCAATAATGTAGATTTACCGACGGA AGGG |
| *DRC7* | DRC7-Bir-cod-MluI-F | AATT**ACGCGT**CTCATTACCCTGCTAATGAATAGATTC |
| | DRC7-Bir-cod-BamHI-R | AATT**GGATCC**CTTTCTATGTAAGGCTGCTAAACG |
| | DRC7-Bir-3UTR-PstI-F | AATT**CTGCAG**CGCTAGATAAAAATATTAGATTGGGCA |
| | DRC7-Bir-3UTR-XhoI-R | AATT**CTCGAG**CTCAGATGATTTACAAATAACTCCTAGTG |
| *DRC9* | DRC9-Bir-cod-MluI-F | AATT**ACGCGT**AGATCTGTGTAGACTGTTTAAAGAAAATCCTGA |
| | DRC9-Bir-cod-BamHI-R | AATT**GGATCC**TTTCTTTTTCTTTTTGCCTTTCTTTCTC |
| | DRC9-Bir-3UTR-PstI-F | AATT**CTGCAG**GTTTGTACAATTTAAAATTAGTATTCAAACAAAGA |
| | DRC9-Bir-3UTR-XhoI-R | AATT**CTCGAG**AAGCTTGAAGTTAAAATCTTCTATCTCAAATTCCATG |
| Overexpression of DRC proteins under the control of MTT1 promoter | | |
| *DRC3* | DRC3-oex-MluI-F | AATT**ACGCGT**TATGTCGAATTACATTAGCAACTACG |
| | DRC3-oex-BamHI-R | AATT**GGATCC**ATTTTCATGATCGGATTCATCATCTC |
| *DRC4A* | DRC4A-oex-MluI-F | AATT**ACGCGT**TATGCCTCCAAAAAAAGCTAAAGG |
| | DRC4A-oex-BamHI-R | AATT**GGATCC**AGAAGAGACTAAACCAGCAGG |
| *DRC4B* | DRC4B-oex-MluI-F | AATT**ACGCGT**TATGTCGAAAGCTGCTAAAGC |
| | DRC4B-oex-BamHI-R | AATT**GGATCC**ATCTGTGTTAGTAGGTACTAAAGGAT |
| *DRC11B* | DRC11B-oex-MluI-F | AATT**ACGCGT**TATGTCTACGAATTTTTATAATTTGCAATGGAAGCAG |
| | DRC11B-oex-BamHI-R | AATT**GGATCC**TTTCTTCTTTTTCTTAGCTTTACCACC |
